# A mobile sex-determining region, male-specific haplotypes, and rearing environment influence age at maturity in Chinook salmon

**DOI:** 10.1101/2020.04.23.056093

**Authors:** Garrett J McKinney, Krista M Nichols, Michael J Ford

**Author notes:** Corresponding Author: Garrett J. McKinney.

## Abstract

Variation in age at maturity is an important contributor to life history and demographic variation within and among species. The optimal age at maturity can vary by sex, and the ability of each sex to evolve towards its fitness optimum depends on the genetic architecture of maturation. Using GWAS of RAD sequencing data, we show that age at maturity in Chinook salmon exhibits sex-specific genetic architecture, with age at maturity in males governed by large (up to 20Mb) male-specific haplotypes. These regions showed no such effect in females. We also provide evidence for translocation of the sex-determining gene between two different chromosomes. This has important implications for sexually antagonistic selection, particularly that sex-linkage of adaptive genes may differ within and among populations based on chromosomal location of the sex-determining gene. Our findings will facilitate research into the genetic causes of shifting demography in Chinook salmon as well as a better understanding of sex-determination in this species and Pacific salmon in general.

## Introduction

Variation in age at maturity is an important contributor to life history and demographic variation within and among species and is often correlated with variation in other phenotypic traits such as differences in size or growth rate (Stearns 1992). Individuals that mature later are often larger which can increase fecundity or competitive advantage for access to mates, increasing reproductive success (Roff 1992). There is a tradeoff however, where later maturation can increase fecundity but at the cost of increased risk of mortality before reproduction (Stearns 1989). This tradeoff might be particularly critical in semelparous species which experience a single reproductive episode before death. Age at maturity is often assumed to be influenced by many genes of small effect; however, recent studies have shown that the genomics of maturation age can be complex with mixed large-effect and polygenic architecture (Barson et al. 2015, Sinclair-Waters et al. 2020). While there are few cases where the genetic architecture of age at maturity is known, the genetic basis of age at maturation has important implications for how populations respond to selection (Kuparinen and Hutchings 2017) and how age diversity can be recovered if lost.

Optimal maturation age commonly varies between sexes, leading to sexually antagonistic selection. In addition, alternative reproductive tactics associated with differences in age or size at maturity are common across taxa and these tactics are often sex-specific (Gross 1996, Emlen 1997, Henson and Warner 1997). As a result, the ability of each sex to evolve towards its fitness optimum can depend on the genetic architecture of maturation age. When genes controlling sexually antagonistic traits are located on autosomes they are exposed to conflicting selection pressures in males and females, preventing selection from acting optimally in either sex (Chippindale et al. 2001). Mechanisms to resolve this sexual conflict include sex-specific phenotypes resulting from the same alleles (Barson et al. 2015, Czorlich et al. 2018), sex-specific gene regulation (Ellegren and Parsch 2007), or mate choice (Albert and Otto 2005). Sexual conflict can also be resolved if the genes in question are located on the sex chromosome (Roberts et al. 2009). Evolutionary theory proposes that the genes controlling sexually antagonistic traits should be over-represented on the sex-chromosomes (Rice 1984); however, empirical studies paint a more complicated picture (see Mank 2009).

Age and size at maturity are important traits in salmon that influence individual fitness, life history variation, population demographics, and fishery characteristics. Older age at maturity is associated with larger size in salmon which can improve reproductive success in females through increased fecundity (Healey and Heard 1984), greater egg size and maternal provisioning to offspring (Nicholas and Hankin 1988), and the ability to dig deeper redds which might be resistant to scouring and superimposition by other females (Berghe and Gross 1984, Weeber et al. 2010). Male salmon exhibit alternative reproductive tactics associated with age at maturity in multiple salmon species (Maekawa and Onozato 1986, Gross 1991, Fleming 1996, Foote et al. 1997). Large dominant males achieve reproductive success by monopolizing access to females, whereas “sneaker” males take up satellite positions and achieve reproductive success by sneaking in among mating pairs to fertilize eggs (Groot and Margolis 1998). In concert with behavioral and size differences, individual sneaker males outcompete dominant males under sperm competition (Vladić et al. 2010, Young et al. 2013) and as a group can sire large portions of offspring in a population (Ford et al. 2015b). Despite the importance of variation in age at maturity, many populations are exhibiting long-term declines in size and age at maturity (Ricker 1981, Lewis et al. 2015, Ohlberger et al. 2018, Losee et al. 2019) that can lead to loss of life history diversity and decreases in population stability. Potential mechanisms for these widespread declines in size and age at maturity include fisheries induced evolution (Sharpe and Hendry 2009), size-selective marine predation (Ohlberger et al. 2019, Seitz et al. 2019), and hatchery breeding and rearing practices (Hankin et al. 2009).

Hatcheries are commonly used to supplement wild salmon stocks; however, an unintended outcome of hatchery rearing practices is that hatchery-reared males often exhibit a shift towards earlier maturation. This has been observed in both Pacific and Atlantic salmon (Larsen et al. 2004, Good and Davidson 2016). Water temperature and feed rations at hatcheries are often optimized for high growth, which in turn promotes early maturation (Larsen et al. 2019); however, hatchery stocks vary in the proportion of males with premature maturation even when raised under identical conditions suggesting genetic differences in susceptibility to early maturation (Spangenberg et al. 2015). Hatchery mating practices, which are often random with respect to size and age, mightalso have inadvertently selected for younger fish (Hankin et al. 2009).

The genetics of age at maturity is still poorly understood in salmonids; however, studies to date appear to show different mechanisms underlying variation in age at maturity among species. In Atlantic salmon, a single gene (VGLL3) explains 39% of the variation in age at maturity in European populations (Barson et al. 2015) but does not appear to influence age at maturity in North American populations (Boulding et al. 2019). In an aquacultural strain of Atlantic salmon, multiple genomic regions, including the VGLL3 gene, explained a total of 78% of the variation in age at maturity (Sinclair-Waters et al. 2020). In Chinook salmon, the specific genes underlying variation in age at maturity are unknown but GWAS has identified SNPs associated with age at maturity on several autosomes (Micheletti and Narum 2018, Waters et al. 2018) and male-specific sex chromosome haplotypes are associated with variation in size and age at maturity in male Chinook salmon from Alaska (McKinney et al. 2019b). Despite the lack of specific knowledge of genes governing age at maturity in most salmon species, studies have consistently shown high heritability for this trait (Gall et al. 1988, Heath et al. 2002, Reed et al. 2018) and QTL/GWAS studies have identified genomic regions associated with age at maturity in multiple species (Moghadam et al. 2007, Haidle et al. 2008, Ayllon et al. 2015). In Chinook salmon, several lines of evidence point to genes on the sex chromosome as strongly influencing age at maturity in this species. This includes sex-linked heritability (Hankin et al. 1993), heritability of male reproductive strategies (Heath et al. 2002), and male-specific haplotypes associated with size and age at maturity in Chinook salmon from Alaska (McKinney et al. 2019b). While the sex chromosome (Ots17) has been strongly implicated in sex-specific age at maturity, genes on other chromosomes have also shown associations (Micheletti and Narum 2018, Waters et al. 2018).

In this study, we examine the genetic basis of age at migration in natural- and hatchery-origin Chinook salmon from the Wenatchee River, Washington, USA. Using RADseq data, we provide evidence for translocation of the sex-determining region among two different chromosomes (Ots17 and Ots18), the first evidence of multiple sex-determining regions in Chinook salmon. The genetic basis of age at maturity varied by sex and by origin. Age at maturity and life-history variation in males were significantly associated with a 15 Mb region of Ots17 that contains male-specific haplotypes; this region showed no association in females. There was a much stronger association between the Ots17 region and age at maturity for fish that spawned and reared in the natural environment compared to those reared in the hatchery environment. Our results have important implications for understanding the causes of long-term demographic shifts in Chinook salmon, such as whether selective predation or fisheries induced evolution is occurring, and provides a foundation to better understand the causes of early maturation in hatcheries.

## Materials and Methods

We examined spring-run Chinook salmon that spawn in the Wenatchee River, a tributary of the Columbia River, east of the Cascade Mountains in western North America. The samples included in this study are a subset of those examined by (Ford et al. 2015a), where the study population and sampling design are detailed. Briefly, mature fish returning to spawn were trapped at a common collection point, Tumwater Dam, below all major spawning areas. At Tumwater Dam, sex, length, weight, and date of sampling were recorded for each fish prior to passing the fish above the dam to continue its spawning migration. Depending on year and location, sex was determined in a variety of ways including external morphology, ultrasound, and observed spawning behavior. Scales were taken from each fish and read for aging. A caudal fin clip was taken and dried on Whatman paper for genetic analysis.

We examined both hatchery- and natural-origin fish, where a hatchery-origin refers to fish whose parents were spawned in a hatchery and natural-origin refers to fish whose parents spawned in the natural stream, regardless of the parents’ ancestry. A hatchery program was established on the Chiwawa River, a major spring-run Chinook spawning tributary of the Wenatchee River, in 1989 to supplement the wild population; this hatchery uses a mixture of natural and hatchery origin fish captured within the watershed each year for broodstock. Similarly, approximately 50%-80% of the natural spawners in a given year are hatchery-origin fish (Ford et al. 2013). The high rates of exchange between the hatchery broodstock and the natural spawning population make this an ‘integrated’ hatchery program with the goal of minimizing genetic divergence between the hatchery and natural groups (Mobrand et al. 2005). Hatchery fish were identified by an adipose fin clip and/or presence of a coded-wire tag. A total of 570 fish returning to the Wenatchee River between 2004 and 2009 were used for RAD sequencing, 205 were natural-origin and 365 were hatchery-origin (Table S1).

Wenatchee spring Chinook salmon exhibit a ‘stream-type’ life-history (Healey 1983) in which the juvenile salmon spend a full year rearing in freshwater after a winter of incubation in the gravel and prior to smolting and migrating to the ocean. The fish then typically spend one to three years in the ocean before returning to spawn at ages ranging from 3 to 5 years-old (Mullan et al. 1992, Ford et al. 2015b). Females exhibit less variance in age at maturity than do males, with most females returning as 4 or 5 year-olds and rarely as 3 year-olds. In contrast, 3 year-old males (also known as ‘jacks’) can make up a substantial portion of the male spawning population. In some years, substantial numbers of males mature precocially, either as parr that do not migrate from the Wenatchee River or as ‘mini-jacks’ that make a short migration to the Columbia River before returning in the same year (as 2 year-olds) to spawn (Harstad et al. 2014, Ford et al. 2015b).

DNA was extracted using the Qiagen DNeasy extraction kit, and sequencing libraries were prepared following the methods of Baird et al. (2008) using *SbfI.* Libraries were sequenced on a HiSeq 2000 or 2500 with single-end 100bp reads; 48 samples were sequence per lane.

RAD sequence data were analyzed using STACKS (V 1.48) (Catchen et al. 2011, Catchen et al. 2013). Default settings were used with the following exceptions: process_radtags: remove reads with an uncalled base (-c), rescue barcodes and radtags by allowing a one base mismatch (-r), discard reads with a low quality score (-q), remove reads marked as failing by Illumina (-filter_illumina) and trim reads to 94 bp length (-t 94), ustacks: bounded SNP model (--model_type bounded) with a maximum error rate of 0.01 (--bound_high 0.01), cstacks: 2 mismatches allowed between loci when building the catalog (-n 2). These settings were used for consistency with previous RADseq analyses of Chinook salmon (McKinney et al. 2016, McKinney et al. 2017a, McKinney et al. 2019a, McKinney et al. 2019b). The --catalog option in cstacks was used to add 10 random samples from this study to the *STACKS* catalog from McKinney et al. (2019b). This allowed the addition of SNPs that might be specific to the Wenatchee population while ensuring consistent locus names between studies.

Quality filters implemented in R scripts were used to identify and remove poor quality and likely uninformative loci and samples. Loci and samples with greater than 30% missing data and loci with less than 1% minor allele frequency (MAF) were removed. Paralogs comprise a substantial portion of the salmon genome but yield unreliable genotypes at read depths typical of RADseq (McKinney et al. 2018). Paralogs were identified using *HDplot* (McKinney et al. 2017b) and removed from further analysis. After paralog removal, we compared genotype data across samples to identify potential duplicate samples. Samples were identified as potentially duplicated if they had greater than 90% identical genotypes for the retained loci.

Positional information for each RADseq locus (RADtag) was obtained by aligning sequences to the Chinook salmon genome (Otsh_v1.0, accession GCA_002872995.1, Christensen et al. 2018) using *bowtie2* (Langmead and Salzberg 2012) with default settings. Loci were assigned positions if they had a full-length (94bp) alignment to the genome with no indels and less than 4 mismatches.

Genome-wide association studies (GWAS) were conducted to identify markers associated with sex and age at maturity. GWAS was conducted using the Genesis package in R (Gogarten et al. 2019) for mixed-model association testing. In all GWAS models, a genetic relationship matrix (GRM) was used to account for overall genetic similarity among individuals due to kinship. Creating the genetic relationship matrix involved three steps. First, a kinship matrix was created using KING (Manichaikul et al. 2010). Second, principle component analysis using PC-Air (Gogarten et al. 2019) was performed on the kinship matrix to generate ancestry representative principle components that describe population structure while accounting for relatedness. Third, the ancestry representative principle components and SNP genotypes were used as input to PCrelate (Gogarten et al. 2019) to obtain pairwise kinship coefficients which were then transformed into the GRM. For each GWAS a null model was fit under the null hypothesis that each SNP has no effect. This model included covariates and the GRM but excluded SNP genotypes. Association tests were then conducted for all SNPs, for each trait, using the fitted null model. For each GWAS, we set the significance threshold at *p*=1.76 × 10^−6^ using Bonferroni correction (α=0.05/# of association tests) to account for multiple testing.

GWAS to identify sex-associated markers was conducted to determine if multiple sex chromosomes exist in this population. The sex chromosome in Chinook salmon has been previously identified as chromosome 17 (Ots17) (Phillips et al. 2013, McKinney et al. 2019b) and the sex-determining gene in Chinook salmon, and most salmonids is *sdY* (Yano et al. 2012, Yano et al. 2013). However, in Atlantic salmon the sex-determining gene *sdY* has translocated to three different chromosomes (Eisbrenner et al. 2014), raising the possibility that *sdY* is present on multiple chromosomes in other salmonid species. A logistic mixed model (Chen et al. 2016) was performed with sex as the dependent variable, coded as 0 (female) and 1 (male), with origin (natural or hatchery) and brood year added as covariates.

GWAS for age at maturity was done separately for males and females and for natural and hatchery origin individuals. Sexes were analyzed separately due to sex-specific differences in distribution of age at maturity and because males and females can differ in the genetic control of age at maturity, for example the VGLL3 gene exhibits sex-specific dominance influencing age at maturity in Atlantic salmon (Barson et al. 2015) and male-specific haplotypes have been associated with variation in size and age at maturity in Chinook salmon from Alaska (McKinney et al. 2019b). Hatchery rearing is also associated with reduced age at maturity but stock-specific effects in similar environments suggest differences in genetic susceptibility to early maturation (Spangenberg et al. 2015). For each sex and origin, a linear mixed model was performed with age at maturity (measured as age at sampling) as the dependent variable and brood year as a covariate.

GWAS for two age-based male life-history traits, jack (age 3) vs non-jack, and precocious (age 2) vs non-precocious, were also performed. These differ from the previous age at maturity GWAS in that these were analyzed as categorical rather than linear traits. These were done because jacks exhibit different spawning behavior than 4 and 5 year old males and because precocious males are a common but undesirable trait seen in hatchery populations. For each GWAS, samples with natural and hatchery origin were analyzed separately because hatcheries have been shown to increase the proportion of jacks. Logistic mixed models were performed with jack (1) vs non-jack (0) or precocious (1) vs non-precocious (0) as the dependent variable and brood year as a covariate.

In addition to GWAS, we also evaluated associations between male-specific haplotypes and age at maturity. Male-specific haplotypes have been previously associated with variation in size and age at maturity in Chinook salmon from Alaska (McKinney et al. 2019b) and we hypothesized that the haplotypes might therefore play a role in variation in size and age at maturity in Wenatchee Chinook salmon. Male-specific haplotypes have been proposed to arise through restricted recombination between the sex chromosomes in Chinook salmon due to male-specific patterns of recombination (McKinney et al. 2019b). Restricted recombination can result in regions of high linkage disequilibrium (LD) spanning several Mb, with different haplotypes characterized by different sets of SNPs in LD. Male-specific haplotypes were identified by conducting network analysis on patterns of LD on the two sex chromosomes identified in this population (Ots17 and Ots18, see results) and by examining other chromosomes for regions of elevated LD that might show sex-specific genotypes. This method has been previously demonstrated to identify and distinguish markers that are part of overlapping genomic features with high LD (McKinney et al. 2020). Within each chromosome, pairwise LD between SNPs was estimated using the *r*^2^ method in *Plink* (V1.9) (Purcell et al. 2007, Chang et al. 2015). Groups of linked SNPs were identified by filtering to marker pairs with *r*^2^ greater than 0.3, then performing network analysis and community detection in R using the igraph package (https://igraph.org/r). Genotypes for groups of linked SNPs were then phased into haplotypes using fastPHASE (Scheet and Stephens 2006). The resulting haplotypes were clustered into haplogroups using heatmap2 (Warnes et al. 2015) in R with the Ward.D clustering algorithm to minimize within group variance.

The association between male-specific haplotypes and age at maturity was tested for significance, and the proportion of variance in age at maturity explained by male-specific haplotypes was estimated using ANOVA (*p* ≤ 0.05) with age as the response variable and haplotype (including unassigned males) as factors. Post-hoc Tukey tests were performed to determine if the average size or age at maturity were significantly different (*p* ≤ 0.05) among male-specific haplotypes. The relationship between male-specific haplotype and size at age was tested for significance (*p* ≤ 0.05) using ANOVA with size (fork length or weight) as the response variable, haplogroup and age as predictor variables, and an interaction between haplotype and age.

## Results

A total of 40,180 SNPs were retained after removing SNPs with more than 30% missing data and less than 1% MAF. Analysis with *HDplot* identified 11,780 SNPs (29%) as paralogs, leaving 28,400 SNPs for the final analysis. Of the retained SNPs, 24,004 (85%) aligned to the genome. A total of 526 samples out of 570 were retained after removing those with more than 30% missing data. Two pairs of apparently duplicated samples were identified with 93% and 94% identical genotypes. All duplicate samples were removed from analysis, leaving 522 samples. The final dataset contained 315 (60%) males and 207 (39%) females (Figure 1, Table S1).

**Figure 1.**
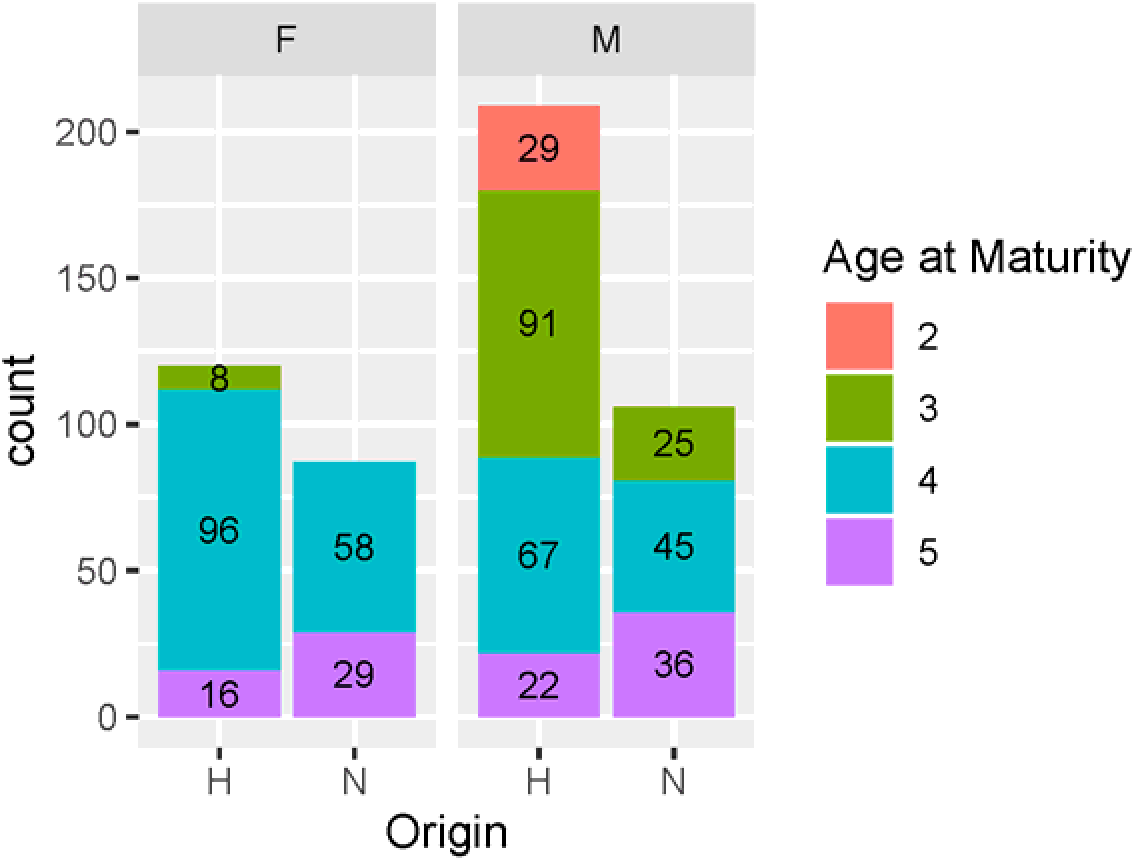
Number of samples retained after quality filtering, reported by sex (male and female), origin (hatchery and natural), and age at maturity.

GWAS of sex resulted in two peaks of association, one on the previously identified sex chromosome (Ots17) (Phillips et al. 2013, McKinney et al. 2019b) and one on Ots18 (Figure 2). A total of 11 SNPs showed significant association after Bonferroni correction (Table S2). Male-specific alleles were identified for nine of these SNPs. On average these alleles occurred in one female (range 0-3) and 41 males (range 36-46).

**Figure 2.**
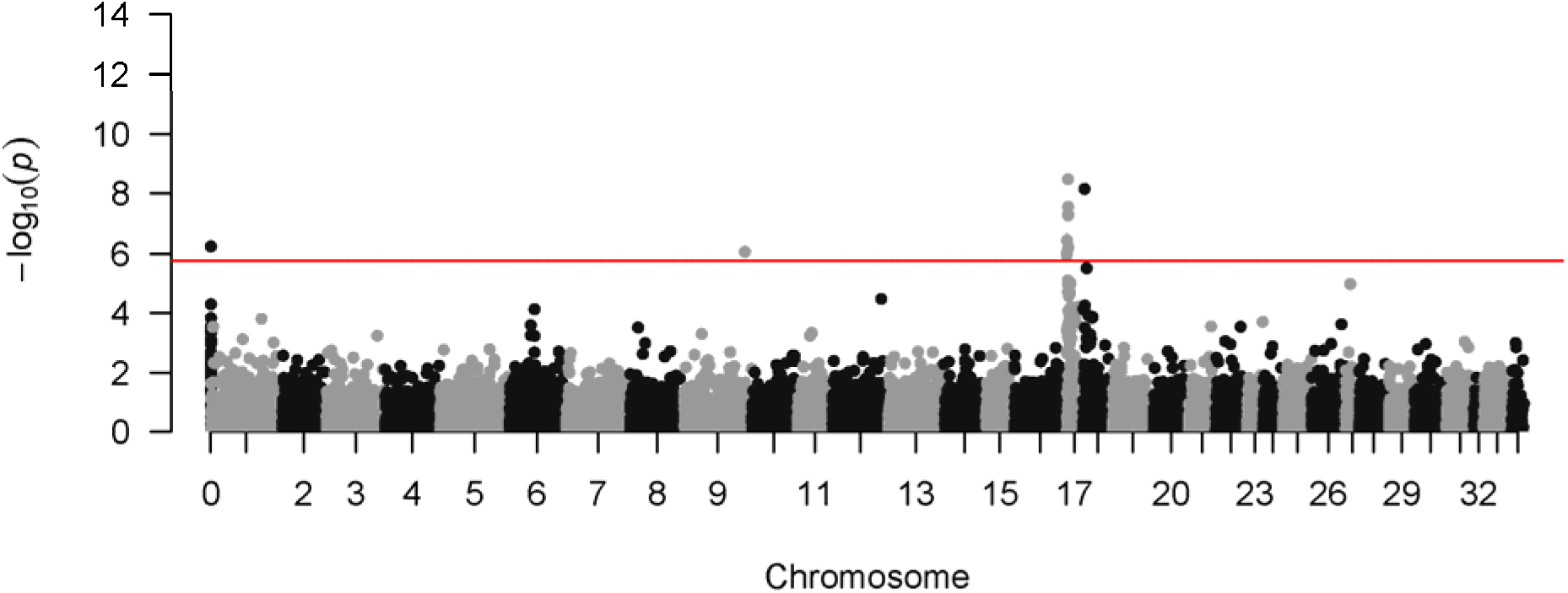
Results for GWAS of sex. Markers not aligned to the genome were assigned to a dummy chromosome (Ots0). Two peaks of association were identified, one on the previously identified sex chromosome (Ots17) and one on Ots18. A single SNP with high association was found on Ots09 and one unmapped SNP was significant. The red line denotes the Bonferroni significance threshold.

GWAS of age at maturity showed different results for males and females of hatchery and natural origin. Natural origin males showed a strong peak of association on Ots17 (Figure 3A). Hatchery males had SNPs significantly associated with age at maturity on multiple chromosomes but not on Ots17 (Figure 3B). A single SNP on Ots03 was associated with age at maturity in natural origin females (Figure 3C) while three were significant in hatchery females, two on Ots18 and one on Ots19 (Table S3, Figure 3D). All SNPs with significant associations in any of the GWAS are reported in Table S3).

**Figure 3.**
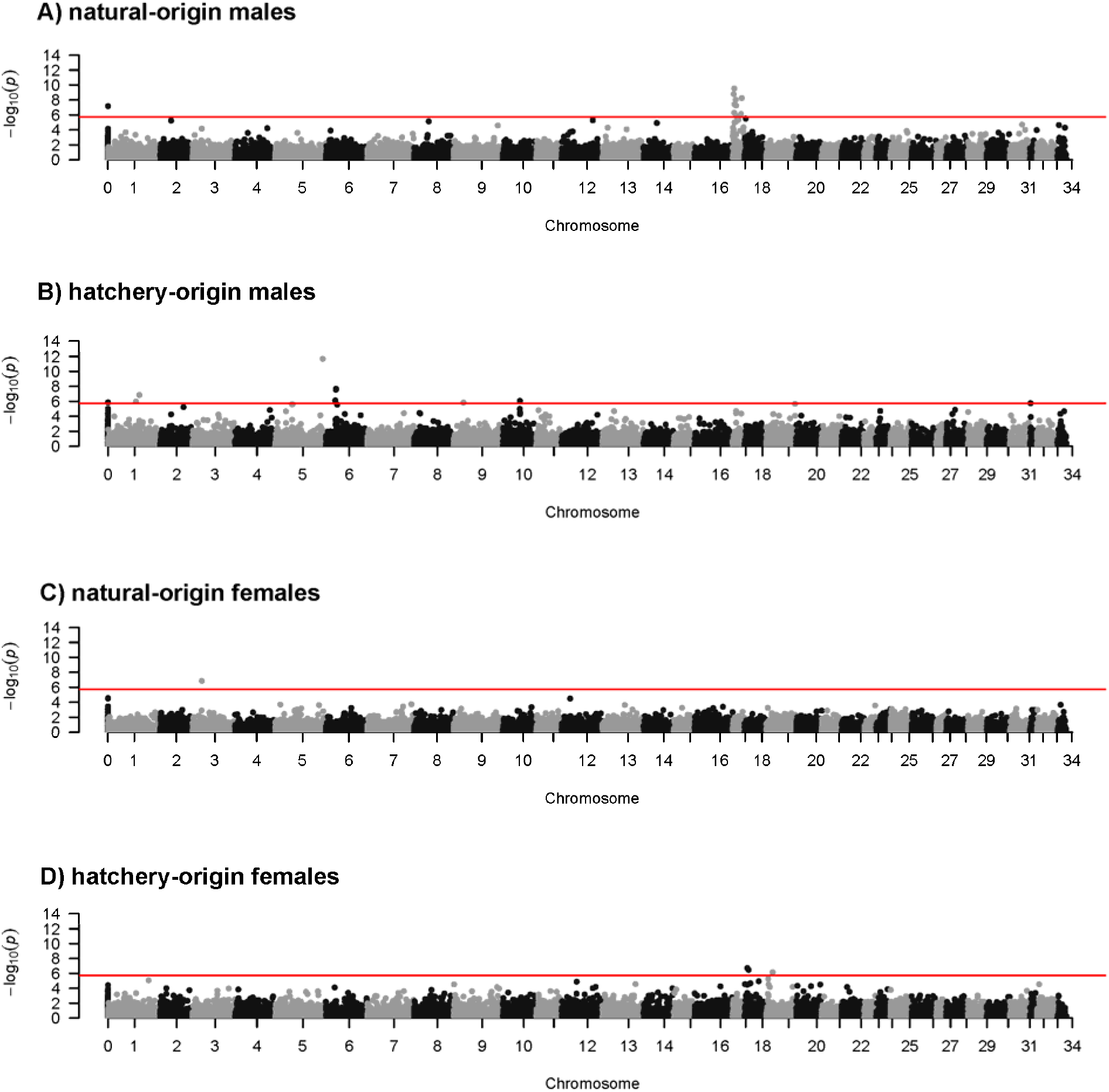
Results of age at maturity GWAS for A) natural-origin males, B) hatchery males, C) natural-origin females, and D) hatchery females. The red line denotes the Bonferroni significance threshold. Markers not aligned to the genome were assigned to a dummy chromosome (Ots0).

GWAS of male life history was conducted for jack (age 3) vs non-jack males and precocious (age 2) vs non-precocious males. When natural and hatchery males were examined together, there was a peak of association with jack life history on Ots17 as well as three other SNPs with significant association (Figure 4A, Table S3). Conducting separate analyses on natural and hatchery males revealed that the peak of association on Ots17 primarily reflected natural males (Figure 4B, Table S3). Hatchery males had a single SNP associated with jack life history on Ots17 as well as four SNPs spread between Ots05, Ots12, and Ots34 (Figure 4C, Table S3). Thirty one SNPs spread among several chromosomes were significantly associated with precocious maturation in hatchery males (Fig 4D, Table S3). These SNPs had low minor allele frequency in non-precocial hatchery males (mean MAF 0.026) and all showed a greater MAF (mean 0.115) in precocial males (Figure S1).

**Figure 4.**
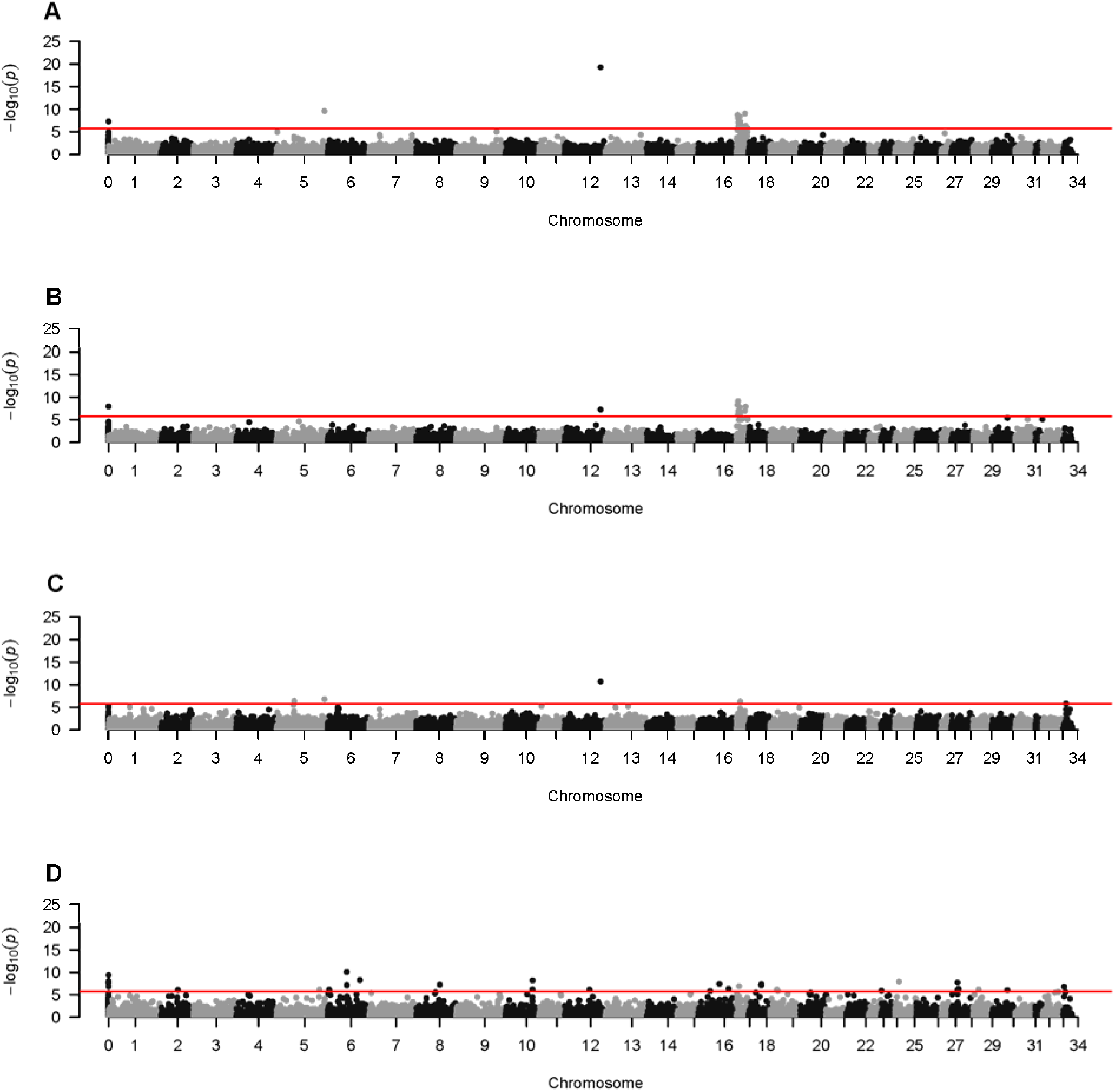
Results of GWAS of male life history. Results for jack vs non-jack males with hatchery-vs natural-origin as a covariate are shown in A. Results for natural-origin jack vs non-jack males only are shown in B. Results for hatchery-origin jack vs non-jack males only are shown in C. Results for hatchery-origin precocious vs non-precocious males are shown in D. The red line denotes the bonferroni significance threshold. Markers not aligned to the genome were assigned to a dummy chromosome (Ots0).

Regions of elevated LD spanning 9 Mb-20 Mb were identified on Ots17, Ots18, and Ots30 (Figure 5). Network analysis identified two sets of linked SNPs on Ots17. One set contained 21 SNPs that spanned 15 Mb. This set contained all the SNPs from Ots17 that were significantly associated with age at maturity in the GWAS. Male-specific alleles at these SNPs formed the Ots17-1 haplogroup (see below). The other set contained 22 linked SNPs spanning 20.5 Mb and contained all SNPs from Ots17 that were significant for the sex GWAS. Male-specific alleles at these SNPs formed the Ots17-2 haplogroup (see below). Two sets of linked SNPs were also found on Ots18, one containing 9 SNPs that spanned 9 Mb and the other containing 35 SNPs that spanned 20 Mb. The SNP on Ots18 that was significantly associated with sex (56111_28) was not part of these LD sets. SNP 56111_28 was filtered out during network analysis because its maximum r^2^ (0.23) fell below the threshold of 0.3 to consider this SNP linked to any other. Finally, two sets of linked SNPs were found on Ots30, one containing 9 SNPs that spanned 20 Mb and one containing 45 SNPs that spanned 33 Mb. No SNPs from Ots30 were associated with sex. The consensus RAD sequence and alleles for all SNPs in these LD blocks are listed in Table S4.

**Figure 5.**
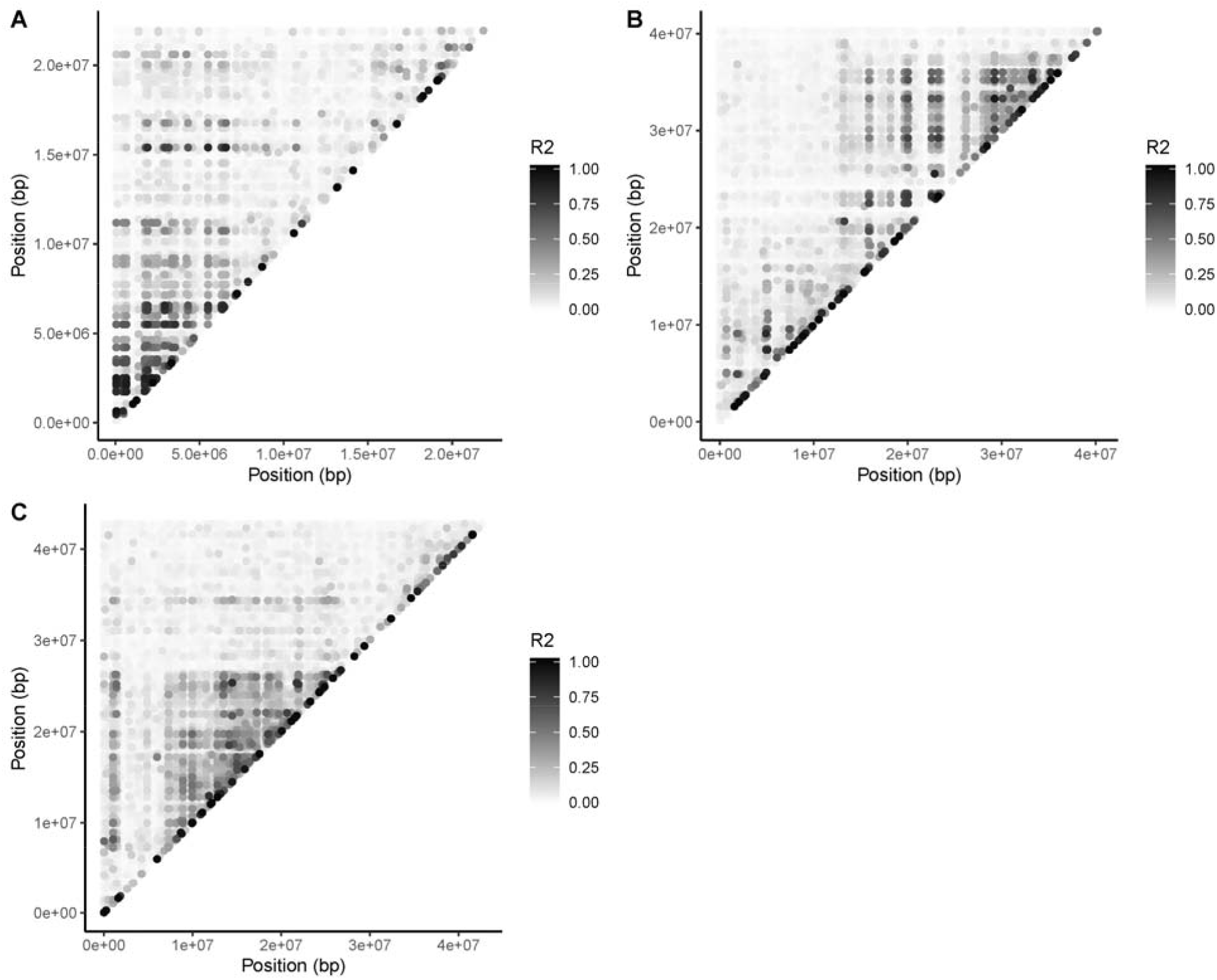
Pairwise LD for chromosomes (A) Ots17, (B) Ots18, and (C) Ots30.

Samples were clustered based on phased haplotypes for high LD SNPs to identify putative male-specific haplogroups. Two clusters of samples were identified on Ots17 that primarily included males (>97%) (Ots17-1, 47 of 48 samples and Ots17-2, 46 of 47) (Figure 6A). Two clusters of samples were also identified on Ots18 (Figure 6B). The Ots18-1 cluster had 90% phenotypic males (56 of 62). The Ots18-2 cluster had only 61% phenotypic males (11 of 18), which is similar to the proportion of males in the full dataset (60%). Six of the Ots18-2 males were also assigned to male-specific haplogroups on Ots17 and two to the Ots18-1 haplogroup. The high number of Ots18-2 males that were also assigned to other male-specific haplogroups along with the high proportion of females assigned to this haplogroup suggested that the LD patterns associated with the Ots18-2 haplogroup were due to a chromosome inversion that is independent of the sex-determining region on Ots18. One cluster of haplotypes was identified on Ots30 (Figure S2), and 77% of the samples in this cluster (20 of 26) were phenotypic males; however, four of the males had been assigned to the Ots18-1 haplogroup. Excluding these samples, the proportion of males decreased to 64%, consistent with the overall sex ratio in this study. This further supported the interpretation that the LD patterns on Ots30 were the result of a chromosome inversion rather than a sex-determining region. In total, 149 of 315 males (47%) were assigned a male-specific haplogroup, 93 to haplogroups on Ots17 and 56 to a haplogroup on Ots18.

**Figure 6.**
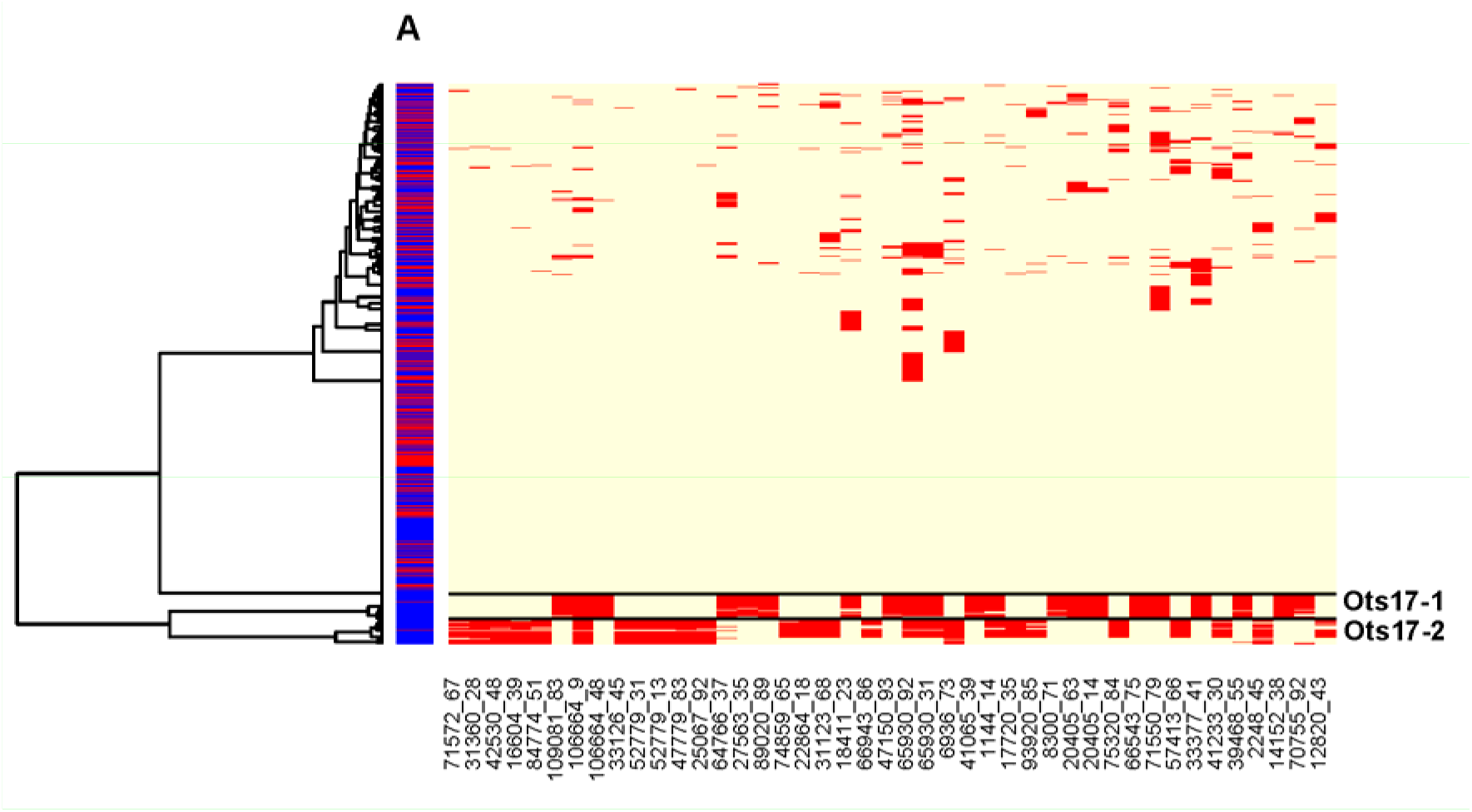
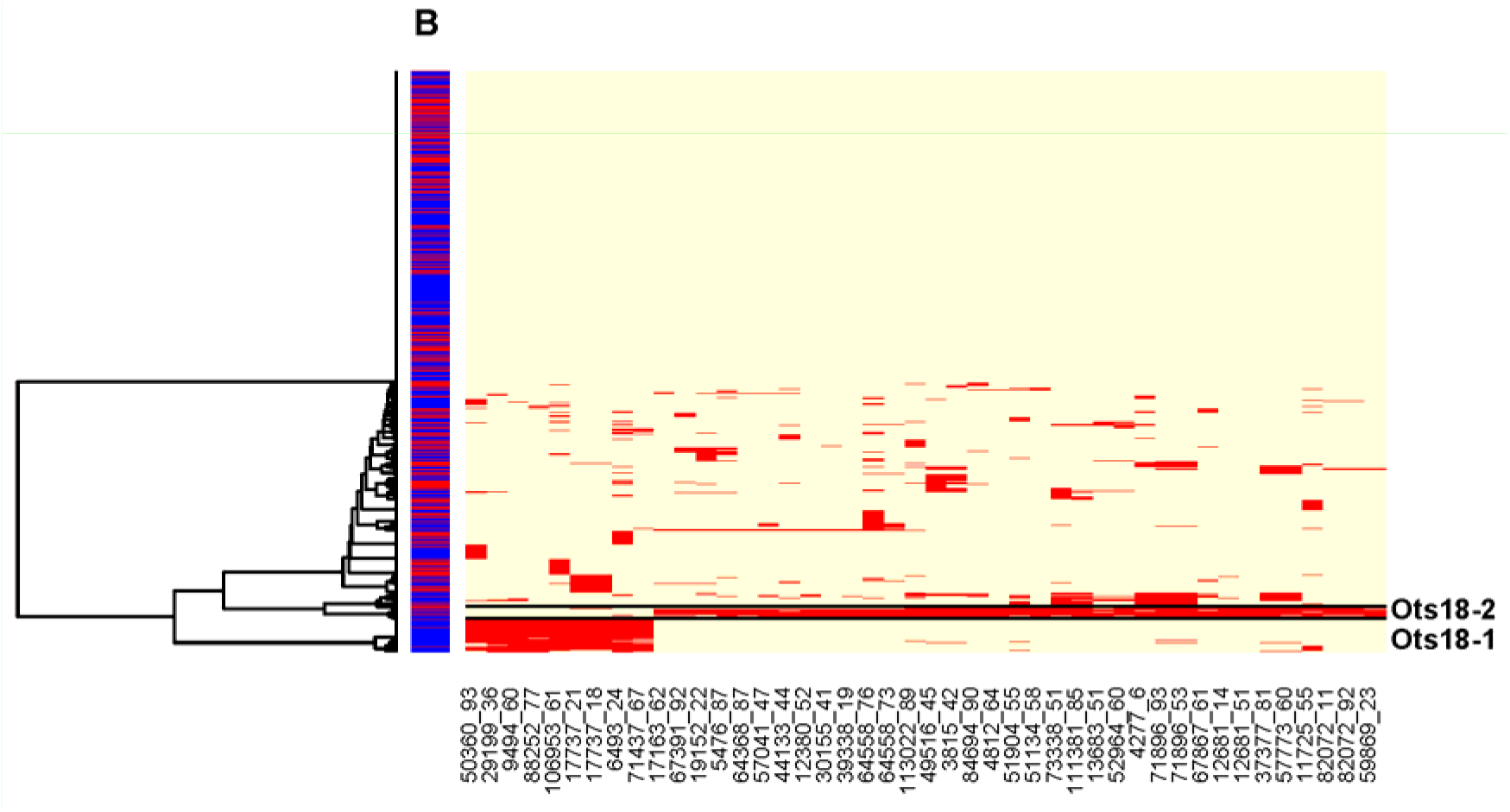
Results of haplotype clustering for sets of high LD loci on A) chromosomes Ots17 and B) Ots18. Individuals are clustered in rows, whereas columns represent loci in order along the chromosomes. Individuals are color coded by sex on the left of the plot, blue for male and red for female. For each SNP, the most frequent allele is in yellow and the least frequent allele is in red. Haplogroups of interest are distinguished by horizontal lines. Clusters Ots17-1, Ots17-2, and Ots18-1 are putative Y-chromosome haplotypes while cluster Ots18-2 is a putative chromosome inversion.

Male-specific haplotypes displayed different distributions of size and age at maturity for male Chinook salmon, and those differences were dependent on hatchery or natural origin (Figure 7). In the natural-origin fish, males with the Ots17-1 haplotype matured at the smallest size (Figure 7A, Figure S3A, Table 1), whereas males with the Ots17-2 and Ots18-1 haplotype matured at the largest size. Males with haplotypes Ots17-1 and Ots17-2 differed on average by approximately 40 cm and 8 kg. Males that could not be assigned to the Ots17-1, Ots17-2, or Ots18-1 haplotypes matured at intermediate sizes. Differences in size at maturity were related to differences in age at maturity (Figure 7B). Males with the Ots17-1 haplotype primarily matured as age three jacks (86%) whereas males with the Ots17-2 and Ots18-1 haplotypes predominantly matured at age five (70%) and none matured younger than age four. The majority of males that could not be assigned a haplotype matured at age four (54%), but 26% matured at age five and 20% at age three. Approximately 48% of the natural-origin jacks (12 of 25) had the Ots17-1 haplotype whereas 53% of the natural-origin, age-5 males (19 of 36) had the Ots17-2 or Ots18-1 haplotypes. Male-specific haplotypes explained 36% of the variance in age at maturity in the natural-origin samples. Hatchery origin males did not show the discrete size distributions for each haplotype that were observed in natural-origin males. Hatchery-origin males with the Ots17-1 haplotype again had the smallest average size and age at maturation. Males with other haplotypes did show an increase in average size or age at maturity relative to the Ots17-1 males but the distributions broadly overlapped. The reduced size at maturity for all haplotypes was driven by a shift towards reduced age at maturity in the hatchery origin fish (Table 1; Figure 7B). Precocious males (age 2) were observed among hatchery-origin fish for all haplotypes but were not observed in natural-origin fish. There was a significant effect of haplotype on length at age in natural-origin male Chinook salmon (*p* < 0.05, Figure S3B). There was a similar trend for weight at age but this was not statistically significant. The influence of haplotype on size at age was most pronounced for fish that matured at age 4 (Figure S3B). For each maturation age observed in natural-origin males (3-5), males with the Ots17-1 haplotype were smallest on average whereas males with the Ots17-2 and Ots18-2 haplotypes were the largest.

**Table 1.** Average length, weight, and age at maturity for hatchery- and natural-origin females and males assigned to each haplogroup.

| Haplogroup | Average Length (cm) |  | Average Weight (kg) |  | Average Age |  |
| --- | --- | --- | --- | --- | --- | --- |
|  | H | N | H | N | H | N |
| Female | 78.80 | 83.50 | 5.36 | 6.27 | 4.07 | 4.33 |
| Ots17_1 | 52.60 | 55.10 | 1.88 | 2.00 | 2.97 | 3.14 |
| Ots17_2 | 68.00 | 97.90 | 5.06 | 10.10 | 3.55 | 4.79 |
| Ots18_1 | 64.60 | 94.20 | 3.66 | 8.80 | 3.35 | 4.62 |
| Unclassified Male | 65.90 | 78.80 | 3.86 | 5.69 | 3.50 | 4.06 |

**Figure 7.**
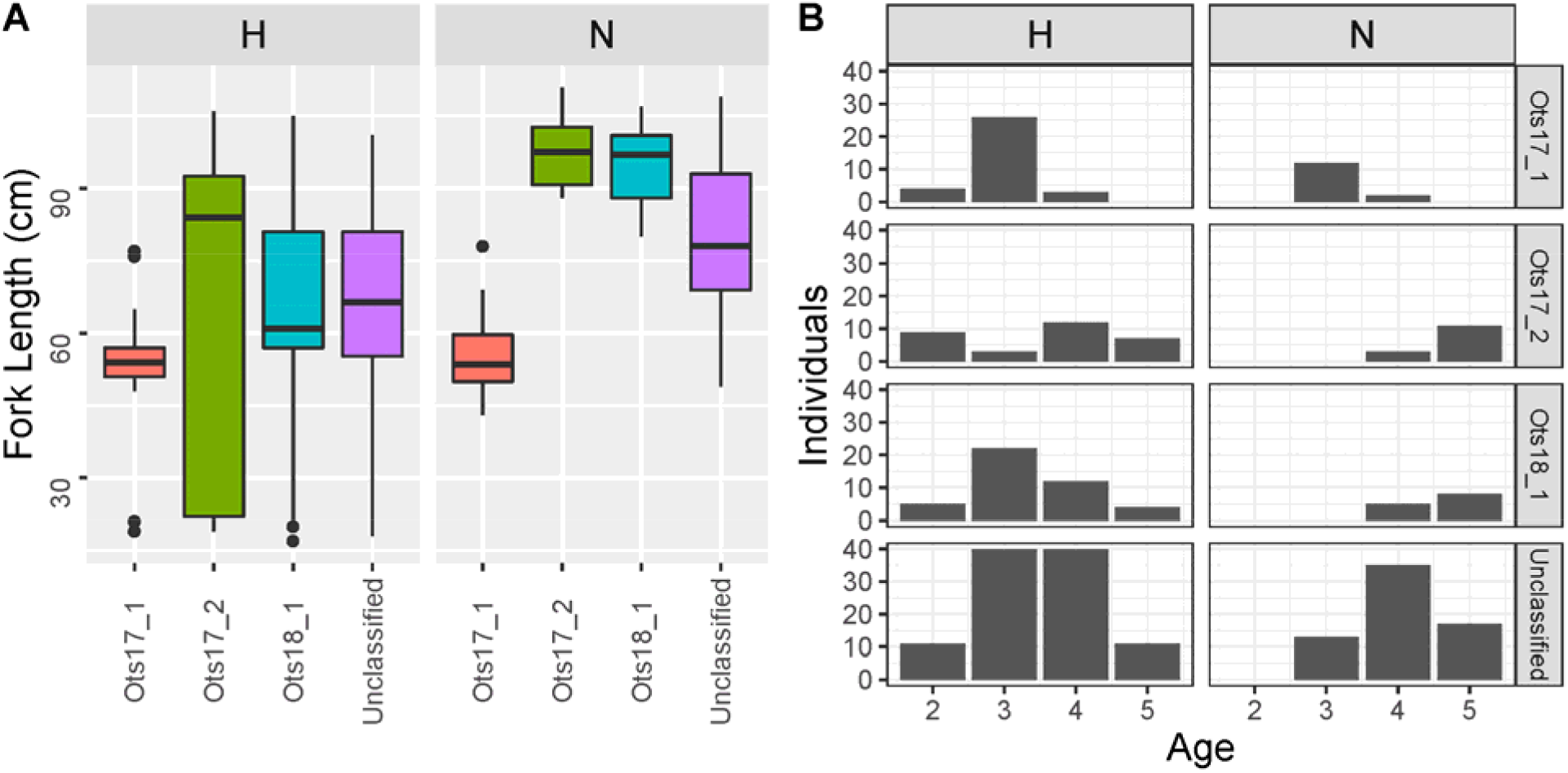
Distributions of A) length at maturity and B) age at maturity for each male-specific haplotype. Significance tests were conducted separately for hatchery- and natural-origin samples. Results of significance tests within each sample origin are included for each panel. Distributions that are significantly different are denoted by different letters, i.e., A is significantly different from B, AB is not significantly different from A or B.

## Discussion

In this study we identified complex genetic control of age at maturity in Chinook salmon, with individual SNPs as well as large male-specific haplotype blocks associated with variation in size and age at maturity. These associations differed by sex and rearing environment, and importantly the sex-linked haplotypes provide a mechanism both for sex-specific selection on age at maturity and for observed sex-specific differences of age at maturity. The SNPs we identified can be used for future examination of context-dependent genetic control of age at maturity. The haplotype-dependent shifts in maturation age in response to the hatchery rearing environment suggests that hatchery rearing conditions are interacting with haplotypes differently than in the natural environment to influence age at maturity. This could be an informative avenue for future research into how to limit early maturation in hatcheries.

Life-history traits such as age at maturity are often assumed to be quantitative and influenced by many genes of small effect. This can lead to inefficient selection when males and females have different fitness optima for maturation age. While few studies have identified a genetic basis to maturation age, it is clear that in some cases age at maturity is influenced by genes of large effect (Yuan et al. 2012) that can exhibit sex-specific effects (Barson et al. 2015). Understanding the genetic architecture of life history traits, even when the causal genes are unknown, can provide important guidance for future research into selection and demographic trends in populations. Size and age at maturity are ecologically and evolutionarily important traits in Chinook salmon that have shown persistent and widespread trends toward younger ages and smaller size over the past four decades (Ricker 1981, Lewis et al. 2015, Ohlberger et al. 2018). Study into the causes of these declines has been complicated by the lack of understanding about the genetic basis of age at maturity in Chinook salmon and how sex and environment might interact with genetics to influence age at maturity.

Size and age at maturity are evolutionarily important traits that often exhibit different fitness optima by sex. One mechanism to resolve this sexual conflict is for causal genetic variants to be located on sex chromosomes so that adaptive alleles can exhibit sex-specific inheritance or expression. This relies on restricted recombination between sex chromosomes. In species without dimorphic sex chromosomes this could be accomplished through heterochiasmy or chromosome inversions. Heterochiasmy is prevalent across taxa on autosomes as well as sex-chromosomes (Lenormand and Dutheil 2005). It is possible that sex-specific haplotypes exist in many species but have not been identified due to lack of genome assemblies or because studies did not examine patterns of LD. Chromosome inversions are also increasingly found to be associated with live-history variation across taxa (Wellenreuther et al. 2019). In salmonids there is strong heterochiasmy in which male recombination is restricted to telomeres (Lien et al. 2011) and large chromosome inversions have been detected in multiple species (Pearse et al. 2019), including on the sex chromosome in chum salmon (McKinney et al. 2020). The large LD-blocks that we identified could be due to strong heterochiasmy or chromosome inversions but we cannot attribute the large LD-blocks to any particular cause with the data available.

In salmon, later maturation is generally favored in females while early maturation in males can reduce the risk of late ocean mortality (Ohlberger et al. 2019, Seitz et al. 2019) or can represent an alternative reproductive tactic with frequency dependent fitness (Berejikian et al. 2010). Male-specific haplotypes linked with *sdY* could resolve sexual conflict by allowing alleles associated with early maturation, such as the Ots17-1 haplotype, to exist in the population without conferring early maturation to females. In concert with this, we found strong genetic influence on age at maturity in male Chinook salmon but few SNPs influencing age at maturity in females. Female Chinook salmon show much less variation in age at maturity than male Chinook salmon. It is not clear from our findings if there less genetic influence on age at maturity in females or if we did not find signals due to recombination between RADseq markers and causal variants.

Despite the *sdY* gene being implicated as the master sex-determining gene in salmonids (Yano et al. 2012, Yano et al. 2013), multiple unrelated chromosome arms have been associated with sex in different salmon species (Phillips et al. 2001, Woram et al. 2003). This suggests that movement of the sex-determining region among species is common. In Atlantic salmon there have also been translocations, and the sex-determining gene has been identified on three different chromosomes (Eisbrenner et al. 2014). Our finding that both Ots17 and Ots18 are linked to sex in Chinook salmon demonstrates that translocation of the sex determining gene has also occurred within this species. In Atlantic salmon, the sex-determining gene *sdY* is flanked by repetitive transposable-like elements that might have facilitated translocation (Lubieniecki et al. 2015); however, it is not known if these same regions flank *sdY* in other species nor whether these repetitive sequences are actually relevant to the movement of *sdY* between chromosomes within and among species. While the only evidence to date of translocations are from Atlantic salmon and Chinook salmon in this study, it is possible that translocations have occurred within other salmonid species but have not yet been identified.

Translocation of the sex-determining region among chromosomes has important implications for the evolutionary potential of populations. Movement of the sex-determining region can cause once-differentiated sex chromosomes to become similar again (Rovatsos et al. 2019). Alternatively, translocation and could enhance adaptation through capture and subsequent sex-linkage of genes (Tennessen et al. 2018). The male-specific haplotypes we identified on Ots17 span overlapping regions of 15 Mb and 20 Mb of the 22 Mb chromosome and contain 481 genes, based on the Chinook salmon genome assembly annotation (Christensen et al. 2018). However, the relative location of *sdY* within this region is unknown because the genome assembly was from a female. These haplotype blocks exclude the telomeric region of Ots17, presumably due to recombination in this region between males and females. Multiple possibilities exist to explain the haplotype influence on age at maturity. The sex-determining region itself might be associated with variation in age at maturity which could explain the later maturation in males with the Ots18-1 haplotype. Age at maturation could also be influenced by genes contained within the broader regions of Ots17 that are part of the Ots17-1 and Ots17-2 haplotype blocks, either through male-specific alleles or through fixed combinations of alleles across multiple genes that are be rare when Ots17 is recombining. Males with the sex-determining region on Ots18 will have two copies of Ots17 that freely recombine in females, losing any male specific alleles that exist on the Ots17 haplotypes and breaking up any co-adapted gene complexes that exist. Variation in the presence and frequency of haplotypes could have important implications for the adaptive potential of populations, particularly for population demography. For example, the observed age distribution varies by haplotype in this study. If the frequency of each haplotype changed this would be expected to shift the overall age distribution. In an extreme case, say fixation of the Ots17-1 or Ots17-2 haplotypes, some maturation ages could be completely lost from the population.

Males with one of three male-specific haplotypes (Ots17-1, Ots17-2, Ots18-1) represented the extremes of maturation age for natural-origin fish in this study (ages 3 and 5). Approximately 53% of the males in this study could not be assigned to one of these haplotypes; these males matured at all age classes but predominantly at age 4. Males with the Ots17-1 haplotype were almost entirely jacks (3 year old males) in both the hatchery and natural-origin populations. The Males that could not be assigned a male-specific haplotype also produced a significant proportion of jacks, particularly in the hatchery. In contrast, the Ots17-2 and Ots18-1 haplotypes produced no jacks, and primarily age 5 males in natural-origin individuals. Jacks are substantially smaller than other male Chinook salmon (Figure 7A, Figure S3A), which results in restricted access to mates when larger dominant males are present. This should reduce fitness relative to larger males; however, jacks can exhibit alternative reproductive tactics where they gain reproductive success by sneaking in among matings rather than guarding nests (Berejikian et al. 2010) and might escape ocean mortality (c.f. Seitz et al. 2019) by returning to spawn at younger ages. Studies have shown frequency dependent fitness of jacks vs dominant males and these alternative life histories likely represent a bet-hedging strategy (Gross 1985, Berejikian et al. 2010). Our results suggest that male-specific haplotypes are linked to life history variation in male Chinook salmon. This is consistent with previous studies showing paternal heritability for age at maturity and life history variation (Hankin et al. 1993, Heath et al. 1994, Heath et al. 2002).

In addition to the haplotype region of Ots17, three SNPs showed significant associations with the jack life history. Most notably, the SNP on Ots12 was the most significantly associated with being a jack (Figure 4). This SNP exhibited unusual genotype patterns with high heterozygosity but one of the homozygous glasses was represented by only one individual. Scatterplots of allele reads revealed three distinct clusters of genotypes that were consistent with elevated ploidy, two of which had been assigned heterozygous genotypes by the Stacks genotyping algorithm (Figure S4). This suggests that the locus was a paralog that was not identified by *HDplot*. Although the original genotype assignments were incorrect, genotypes were not assigned because the allele ratios did not fit tetraploid expectations either. It is unclear whether a null allele contributes to this pattern or if there is copy number variation at this locus. While paralogs are typically discarded in RADseq data due to issues with genotyping, recent work has identified copy number variation associated with variance in sea surface temperature, suggesting adaptive differences (Dorant et al. 2020).

Hatchery males exhibited earlier maturation for all haplotypes relative to the natural-origin population (Figure 7B). Hatchery rearing influenced fish with alternative haplotypes differently, with the Ots17-1 haplotype showing little change in maturation age, whereas fish with the Ots17-2 and Ots18-1 haplotypes matured an average of a year earlier in the hatchery (Table 1). Natural-origin males with the Ots17-2 and Ots18-1 haplotypes matured only at age 4 and 5, but these haplotypes were found in all age classes in the hatchery-origin fish. Precocious males were only observed in the hatchery but were represented by all haplotypes. This, along with the broad distribution of significant SNPs throughout the genome, suggests that the precocious male phenotype is controlled by many genes of small effect combined with a large environmental influence. It is likely that precocial maturation was triggered by hatchery rearing conditions which tend to favor rapid growth and have been demonstrated to cause early maturation in salmon (Larsen et al. 2006). It is unclear if the different shifts in maturation represent a true difference in susceptibility among haplotypes to early maturation under hatchery conditions, or if it is a result of a lower limit on maturation age. Paradoxically, if large old fish are being selected against in the wild, the hatchery might confer a protective effect as the haplotypes associated with the oldest and largest fish in the wild show the largest shift towards early maturation in the hatchery. Maturing at a younger age might allow these fish to escape ocean mortality and pass late maturation genes on to future generations. Our results suggest also that proposals to use selective breeding in hatcheries to counter declining trends in salmon age and size (Hankin et al. 2009) might be more effective if such selection occurs directly on the Ots17 and Ots18 haplotypes rather than on size itself because larger size associated with haplotype is only expressed in wild fish. Though more work is clearly needed, new insight from the current study into genetic architecture of maturation will be informative for future research into mitigating undesirable early maturation in many hatcheries.

Male-specific haplotypes showed significant differences in length at age and trends of different weight at age in our study that have important implications for observed changes in population demographics. Previous studies have shown that in addition to declines in size and age at maturity, size at age has also been decreasing over the past several decades in many Chinook salmon populations, particularly for older fish (Lewis et al. 2015, Ohlberger et al. 2018, Ohlberger et al. 2019). This change is concerning because Chinook salmon are an important resource for fisheries and marine predators. Salmon fisheries are typically managed with limits on number of fish caught rather than biomass, so smaller fish generally mean less profit for fishermen. Similarly, smaller fish mean marine predators would need to expend more energy hunting to achieve the same number of calories. This is particularly relevant to southern resident killer whales which preferentially feed on large Chinook salmon and increasing populations of killer whales have been hypothesized to be a source of natural selection driving declining trends in Chinook salmon size (Ford and Ellis 2006, Ohlberger et al. 2019). Our results suggest that selection that increases frequency of haplotypes associated with younger age at maturity could also result in reduced size at age. A wider survey of spatial and temporal trends in the frequencies of age-associated male-specific haplotypes would be helpful to further elucidate the causes of these trends and the potential for their reversal.

This study has two important implications for how the genetic basis of maturation is interpreted in prior and future studies. First, studies examining the genetics of age at maturity in salmon often raise fish in hatcheries or under hatchery-like conditions. Our results demonstrate that hatchery rearing conditions obscure the relationship between genotype and phenotype compared to that found in natural conditions. The results of studies of age at maturity might therefore not be transferable across rearing environments. Second, in combination with previous results, it is clear that male-specific haplotypes not only vary in frequency but also identity across the Chinook salmon range. In each case these results are based on likely neutral SNPs that are in LD with causal variants so it is difficult to say whether the genetic basis of male age at maturity differs among populations or if we are observing differences in the surrounding neutral evolutionary history. The Ots17-1 haplotype was not identified in in a previous study of Chinook salmon from Alaska despite extensive sampling, suggesting it might be regionally restricted (McKinney et al. 2019b). There was also no evidence of a sex determining region on Ots18 in Alaskan Chinook. The Ots17-2 haplotype shares a number of SNPs that characterize the AK Y4 haplotype in Alaska, suggesting a common evolutionary origin. This haplotype was also associated with the largest fish in Alaska and the Wenatchee River, suggesting a conserved genetic basis for older age at maturity among these haplotypes. Two previous studies of Chinook salmon failed to find a signal of age at maturity on the Ots17 (Micheletti and Narum 2018, Waters et al. 2018). It is possible that these populations had little or no male-specific haplotype variation to detect; however, it is also possible that pooling samples by age class (e.g. Micheletti and Narum 2018) could have masked signals of male-specific haplotypes. If multiple haplotypes are present, but each at low frequency, there may not be enough individuals with haplotype-specific alleles to reach significance in a GWAS. Even in this study, only one of the male haplotypes contained SNPs significantly associated with age at maturity in the GWAS. We were only able to show the significant association between all haplotypes and age at maturity after the haplotypes were identified. Further study into these male-specific haplotypes, including whole-genome resequencing, are needed to better understand the origin of these haplotypes, heterochiasmy or inversions, and to identify the causal variants underlying phenotypic differences between males with different haplotypes.

## Conclusion

Using GWAS, we found a genomic region strongly associated with variation in male age and size at maturity in Chinook salmon from the Wenatchee River. This region was characterized by multiple male-specific haplotypes that are associated with size and age at maturity. Hatchery origin fish showed shifts towards earlier maturation that were haplotype-specific, suggesting differential genetic susceptibility to early maturation. Male-specific Haplotypes identified in this study included two novel haplotypes and one haplotype that is genetically similar to a male-specific haplotype previously identified in Alaska. Those differences and similarities show that although substantial variation for male-specific haplotypes exists across the species range, there are also related haplotypes that show broad geographic distribution. This mixed result suggests both evolutionary conservation and potential differentiation in the genetic basis of male age at maturity throughout the Chinook salmon range. Our results also provide a mechanism both for resolving sexual conflict in age at maturity in Chinook salmon and for the development of alternative male reproductive tactics. These findings are a significant advance in the understanding of the genomics of age at maturity in salmon and will provide a foundation for further work into the evolution of life history in this and other species.

## Acknowledgements

We would like to thank field collectors for obtaining genetic samples and collecting phenotypic data and Sharon Howard for preparing sequencing libraries. We would also like to thank the Columbia River Inter-tribal Fisheries Commission for sequencing three of the RADseq libraries. Garrett McKinney was supported by a National Research Council postdoctoral fellowship.

---

**Table S1.**
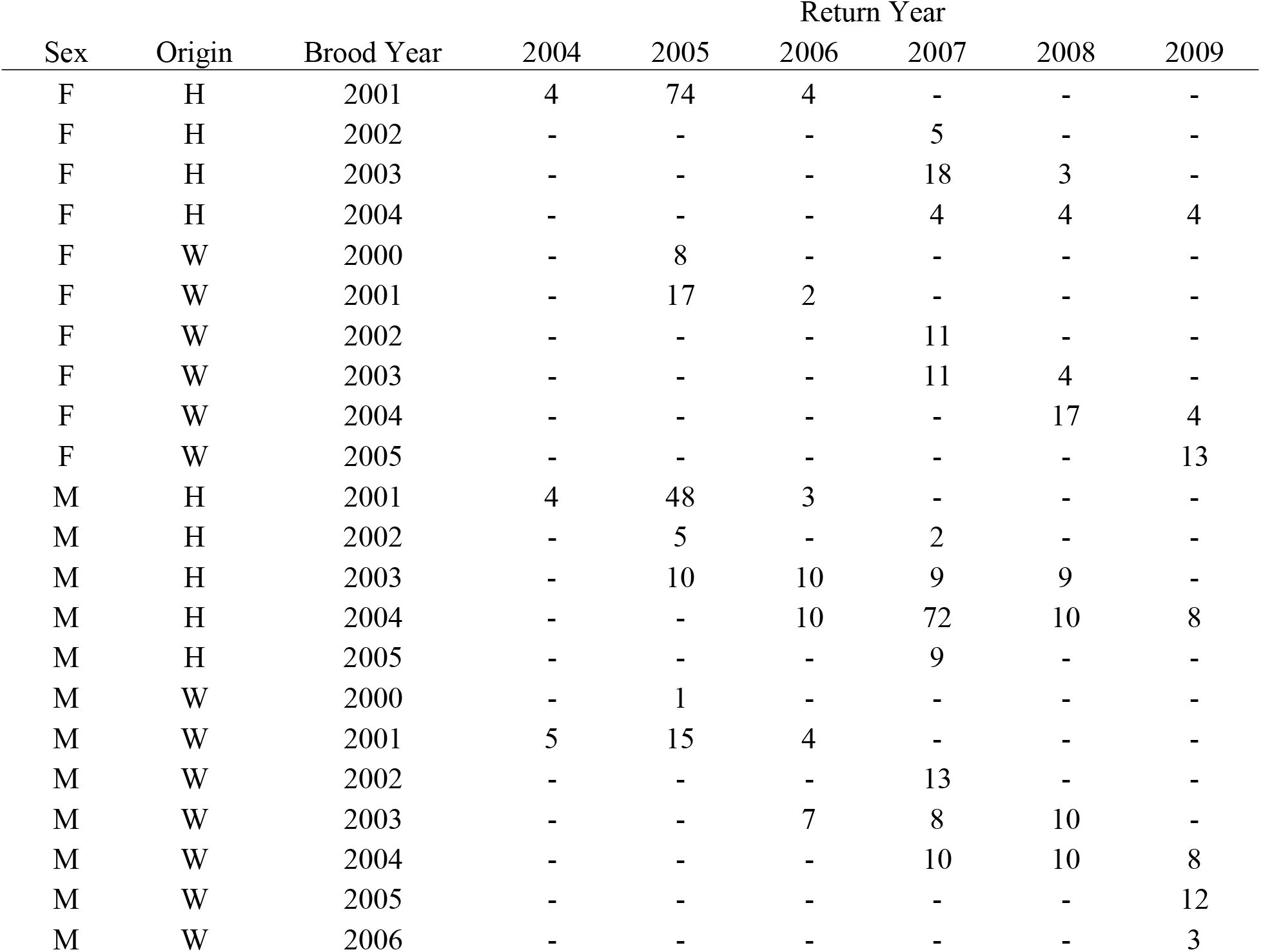
Number of fish retained after quality filtering broken down by sex, origin, brood year, and return year.

**Table S2.**
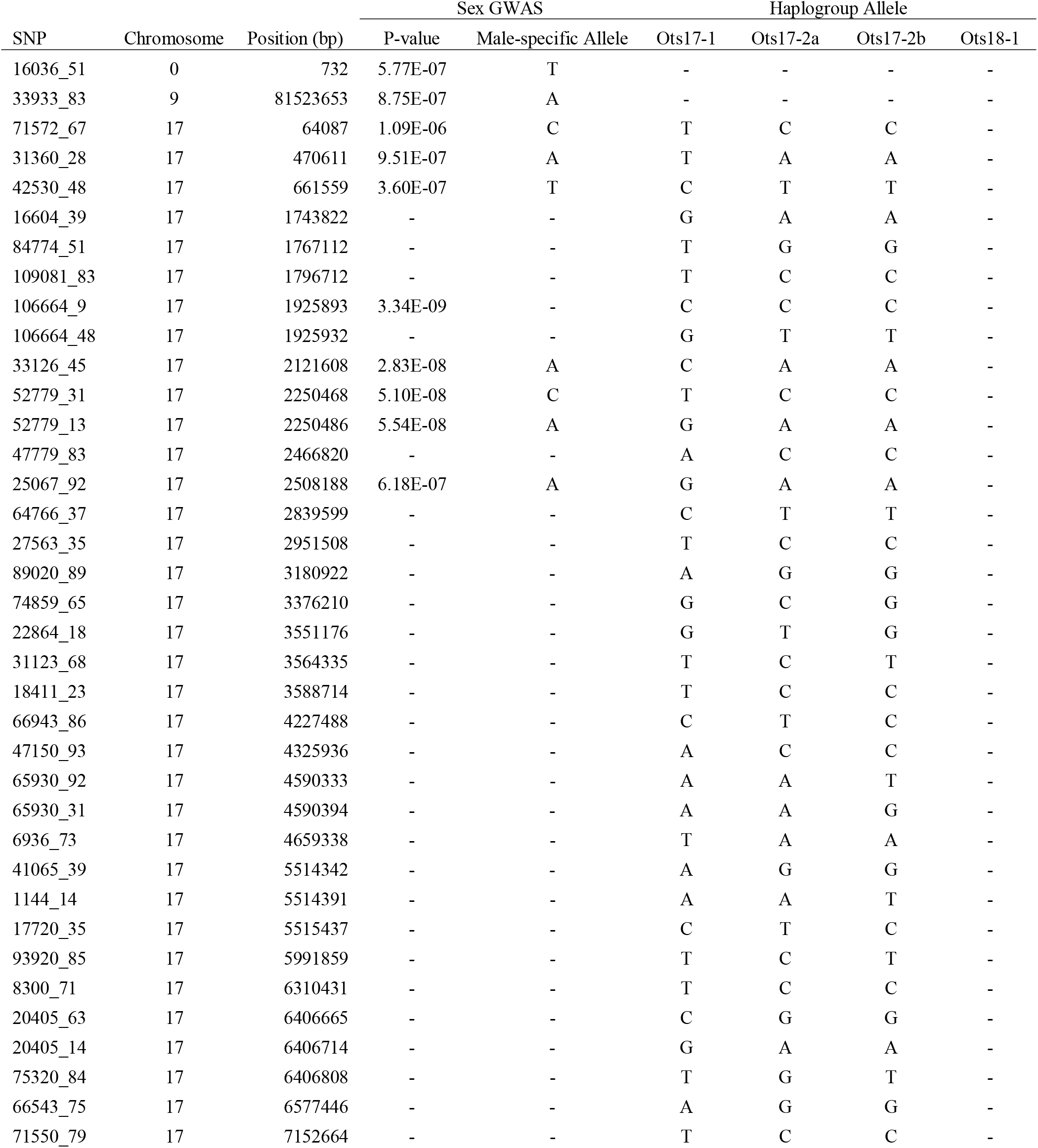
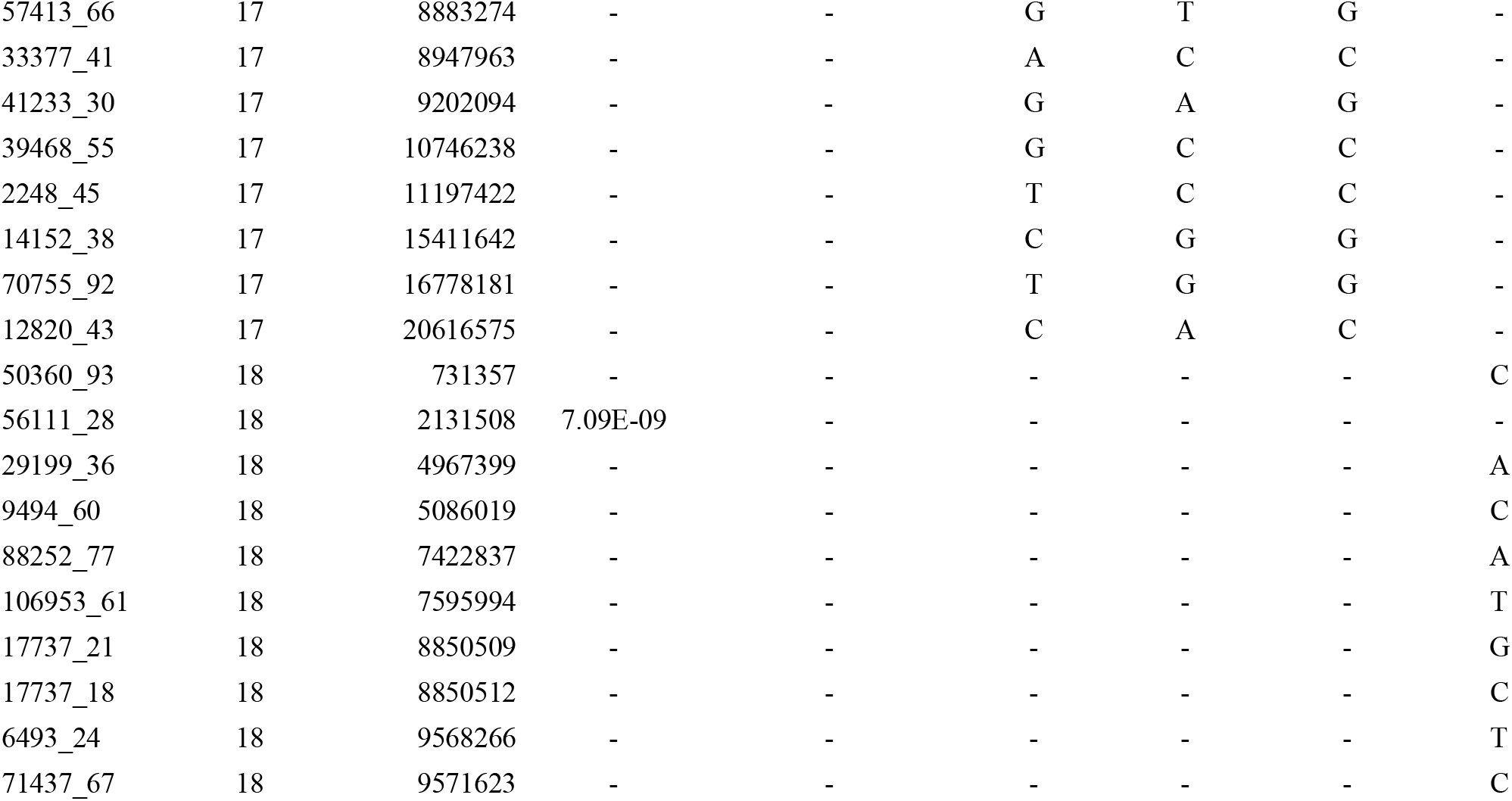
SNPs significantly associated with sex. Putative sex-diagnostic alleles are listed in the Male-specific Allele column. For SNPs that were part of each haplogroup, the alleles associated with that haplogroup are listed. Markers not aligned to the genome were assigned to a dummy chromosome (0) and given arbitrary sequential positions.

| SNP | Chromosome | Position (bp) | Sex GWAS |  | Haplogroup Allele |  |  |  |
| --- | --- | --- | --- | --- | --- | --- | --- | --- |
|  |  |  | P-value | Male-specific Allele | Ots17-1 | Ots17-2a | Ots17-2b | Ots18-1 |
| 16036_51 | 0 | 732 | 5.77E-07 | T | - | - | - | - |
| 33933_83 | 9 | 81523653 | 8.75E-07 | A | - | - | - | - |
| 71572_67 | 17 | 64087 | 1.09E-06 | C | T | C | C | - |
| 31360_28 | 17 | 470611 | 9.51E-07 | A | T | A | A | - |
| 42530_48 | 17 | 661559 | 3.60E-07 | T | C | T | T | - |
| 16604_39 | 17 | 1743822 | - | - | G | A | A | - |
| 84774_51 | 17 | 1767112 | - | - | T | G | G | - |
| 109081_83 | 17 | 1796712 | - | - | T | C | C | - |
| 106664_9 | 17 | 1925893 | 3.34E-09 | - | C | C | C | - |
| 106664_48 | 17 | 1925932 | - | - | G | T | T | - |
| 33126_45 | 17 | 2121608 | 2.83E-08 | A | C | A | A | - |
| 52779_31 | 17 | 2250468 | 5.10E-08 | C | T | C | C | - |
| 52779_13 | 17 | 2250486 | 5.54E-08 | A | G | A | A | - |
| 47779_83 | 17 | 2466820 | - | - | A | C | C | - |
| 25067_92 | 17 | 2508188 | 6.18E-07 | A | G | A | A | - |
| 64766_37 | 17 | 2839599 | - | - | C | T | T | - |
| 27563_35 | 17 | 2951508 | - | - | T | C | C | - |
| 89020_89 | 17 | 3180922 | - | - | A | G | G | - |
| 74859_65 | 17 | 3376210 | - | - | G | C | G | - |
| 22864_18 | 17 | 3551176 | - | - | G | T | G | - |
| 31123_68 | 17 | 3564335 | - | - | T | C | T | - |
| 18411_23 | 17 | 3588714 | - | - | T | C | C | - |
| 66943_86 | 17 | 4227488 | - | - | C | T | C | - |
| 47150_93 | 17 | 4325936 | - | - | A | C | C | - |
| 65930_92 | 17 | 4590333 | - | - | A | A | T | - |
| 65930_31 | 17 | 4590394 | - | - | A | A | G | - |
| 6936_73 | 17 | 4659338 | - | - | T | A | A | - |
| 41065_39 | 17 | 5514342 | - | - | A | G | G | - |
| 1144_14 | 17 | 5514391 | - | - | A | A | T | - |
| 17720_35 | 17 | 5515437 | - | - | C | T | C | - |
| 93920_85 | 17 | 5991859 | - | - | T | C | T | - |
| 8300_71 | 17 | 6310431 | - | - | T | C | C | - |
| 20405_63 | 17 | 6406665 | - | - | C | G | G | - |
| 20405_14 | 17 | 6406714 | - | - | G | A | A | - |
| 75320_84 | 17 | 6406808 | - | - | T | G | T | - |
| 66543_75 | 17 | 6577446 | - | - | A | G | G | - |
| 71550_79 | 17 | 7152664 | - | - | T | C | C | - |
| 57413_66 | 17 | 8883274 | - | - | G | T | G | - |
| 33377_41 | 17 | 8947963 | - | - | A | C | C | - |
| 41233_30 | 17 | 9202094 | - | - | G | A | G | - |
| 39468_55 | 17 | 10746238 | - | - | G | C | C | - |
| 2248_45 | 17 | 11197422 | - | - | T | C | C | - |
| 14152_38 | 17 | 15411642 | - | - | C | G | G | - |
| 70755_92 | 17 | 16778181 | - | - | T | G | G | - |
| 12820_43 | 17 | 20616575 | - | - | C | A | C | - |
| 50360_93 | 18 | 731357 | - | - | - | - | - | C |
| 56111_28 | 18 | 2131508 | 7.09E-09 | - | - | - | - | - |
| 29199_36 | 18 | 4967399 | - | - | - | - | - | A |
| 9494_60 | 18 | 5086019 | - | - | - | - | - | C |
| 88252_77 | 18 | 7422837 | - | - | - | - | - | A |
| 106953_61 | 18 | 7595994 | - | - | - | - | - | T |
| 17737_21 | 18 | 8850509 | - | - | - | - | - | G |
| 17737_18 | 18 | 8850512 | - | - | - | - | - | C |
| 6493_24 | 18 | 9568266 | - | - | - | - | - | T |
| 71437_67 | 18 | 9571623 | - | - | - | - | - | C |

**Table S3.**
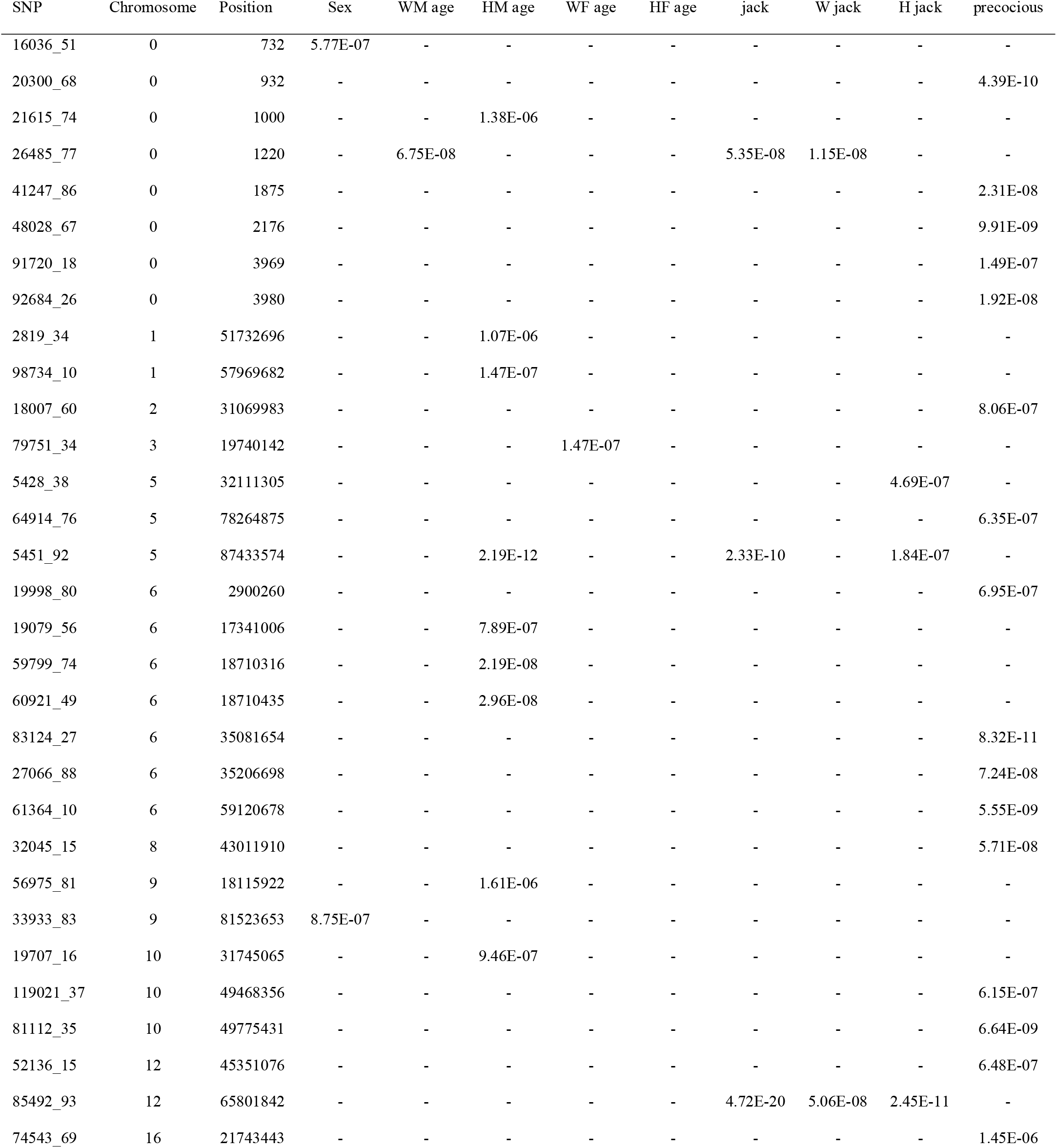
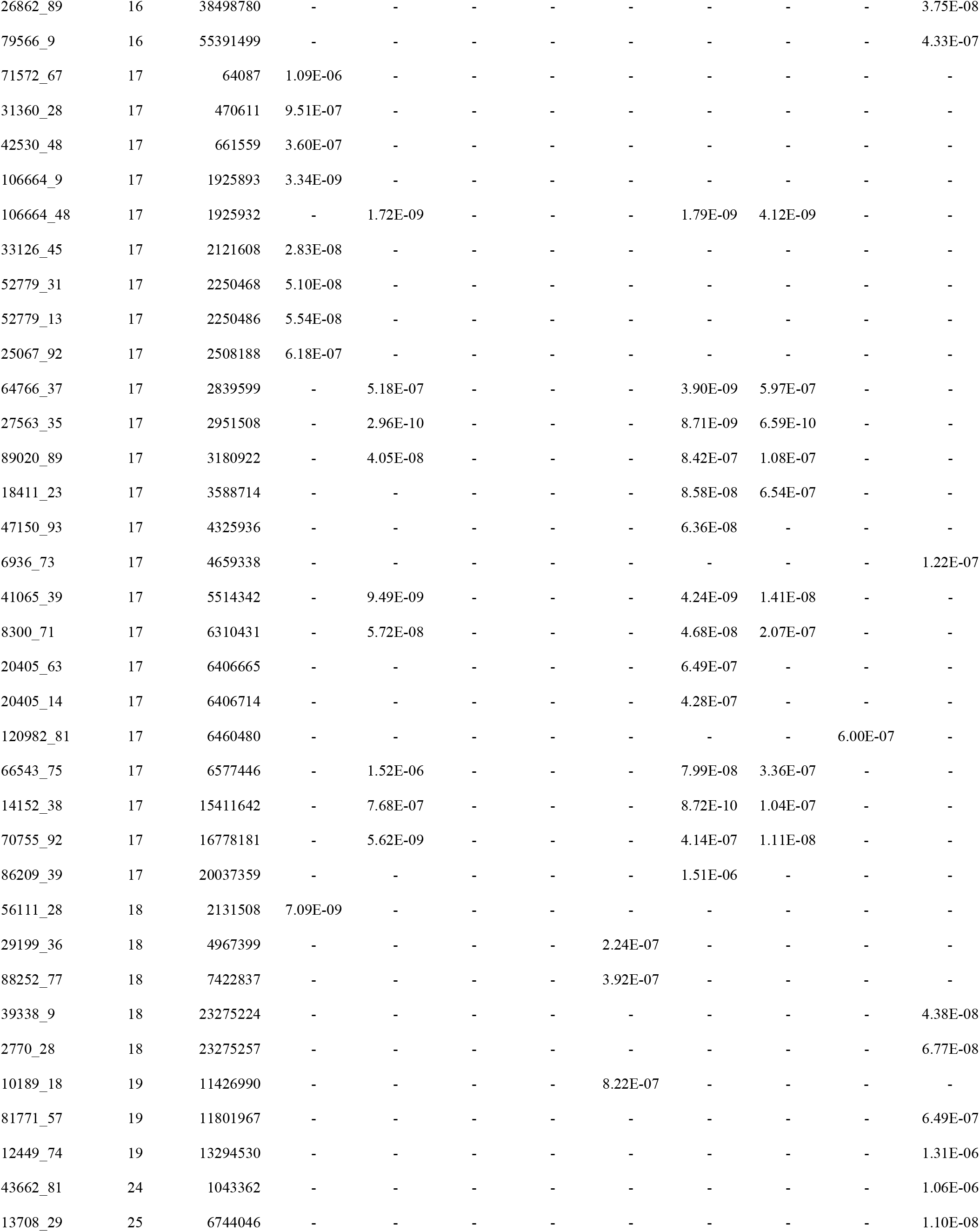
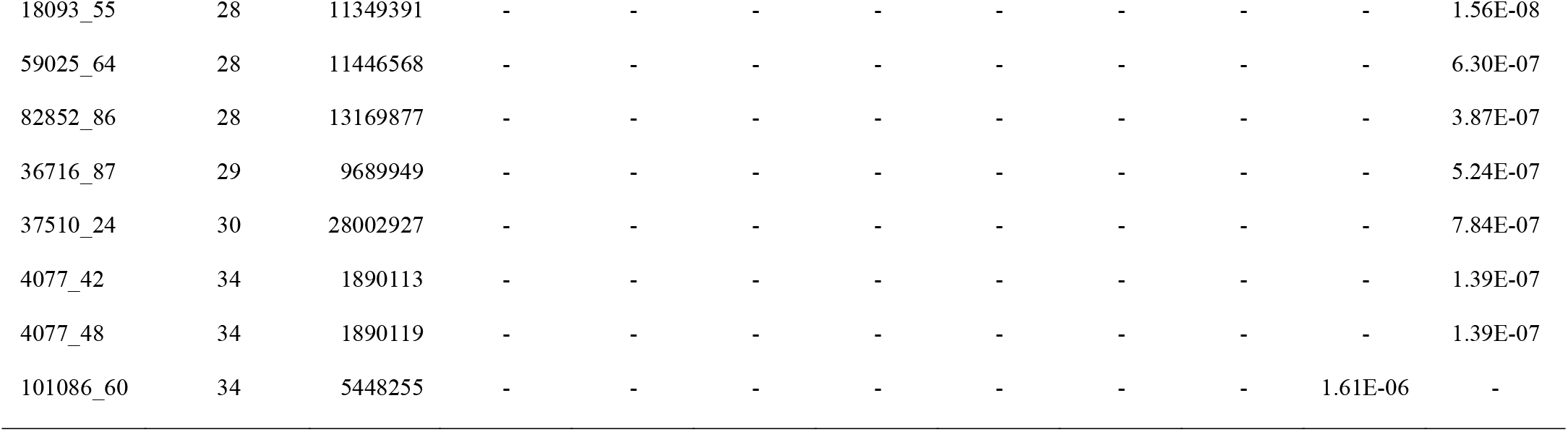
Location of significant SNPs for each GWAS. For traits where samples were examined separately by origin and sex, N and H denote natural- and hatchery-origin and M and F denote male and female. P-values are listed for significant SNPs only. Markers not aligned to the genome were assigned to a dummy chromosome (0) with arbitrary sequential positions.

**Figure S1.**
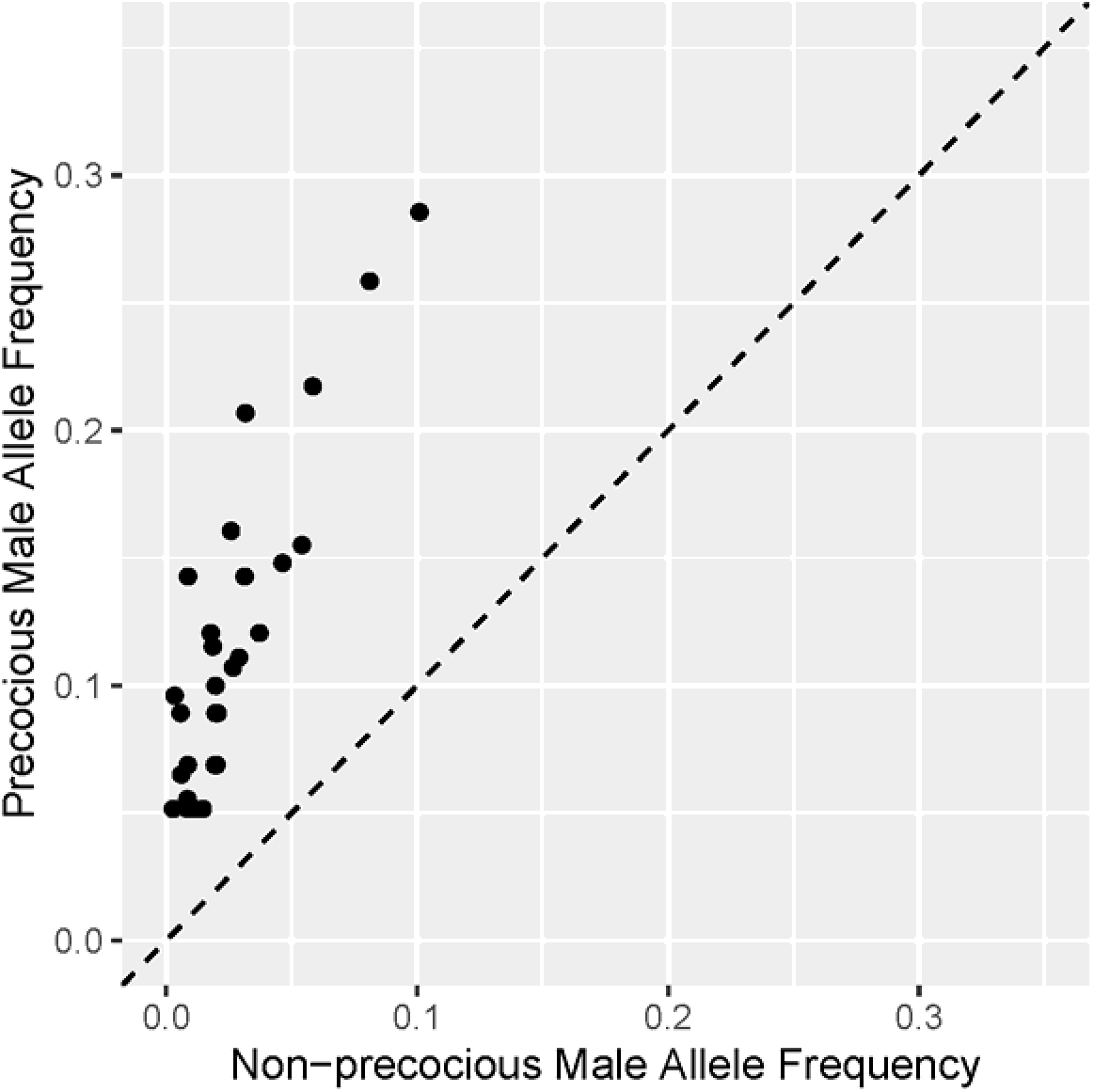
Scatterplot of minor allele frequencies in precocious and non-precocious hatchery males for SNPs significantly associated with precocial maturation. SNPs above the dashed line had a higher minor allele frequency in precocious relative to non-precocious males.

**Figure S2.**
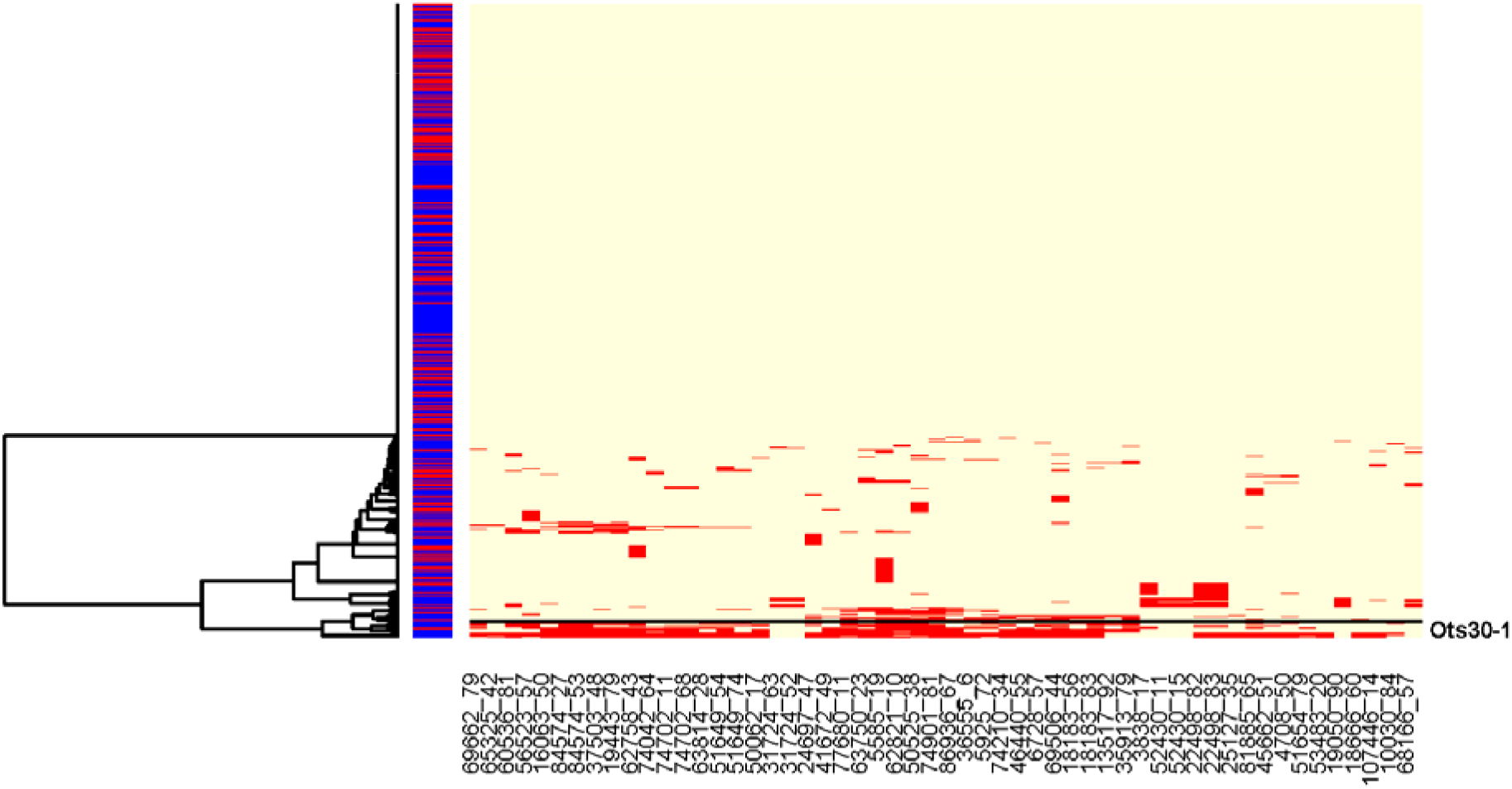
Results of haplotype clustering for sets of high LD loci on Ots30. Samples are clustered in rows while loci are in order along the chromosomes in columns. Samples are color coded by sex on the left of the plot, blue for male and red for female. For each SNP, the most frequent allele is in yellow and the least frequent allele is in red. Haplogroups of interest are distinguished by horizontal lines. Cluster Ots30-1 is a putative chromosome inversion.

**Figure S3.**
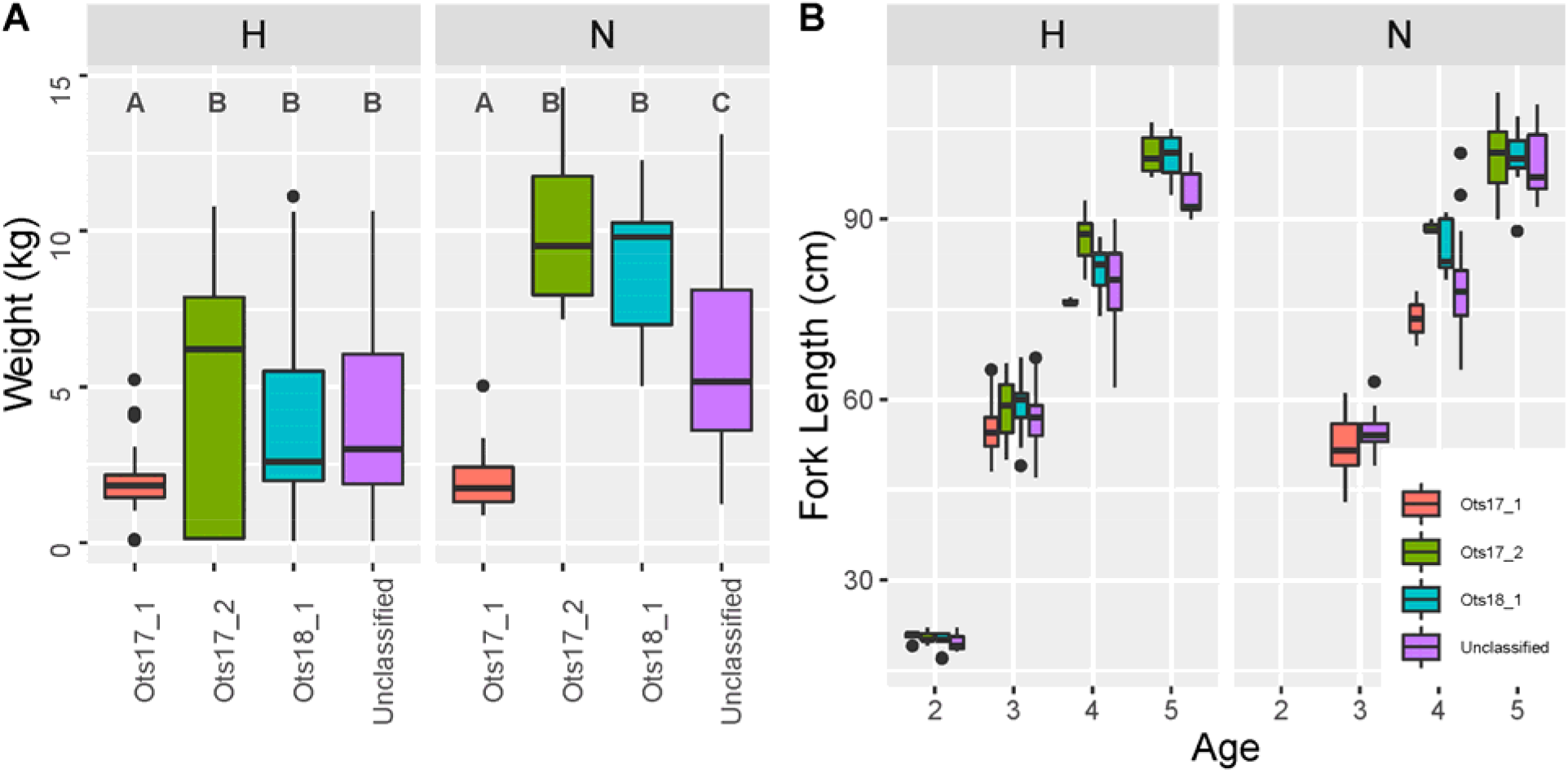
Distributions of A) weight at maturity and B) length at age for each male-specific haplotype. Significance tests were conducted separately for hatchery- and natural-origin samples. Results of significance tests within each sample origin are included for each panel. Distributions that are significantly different are denoted by different letters, i.e. A is significantly different from B.

**Figure S4.**
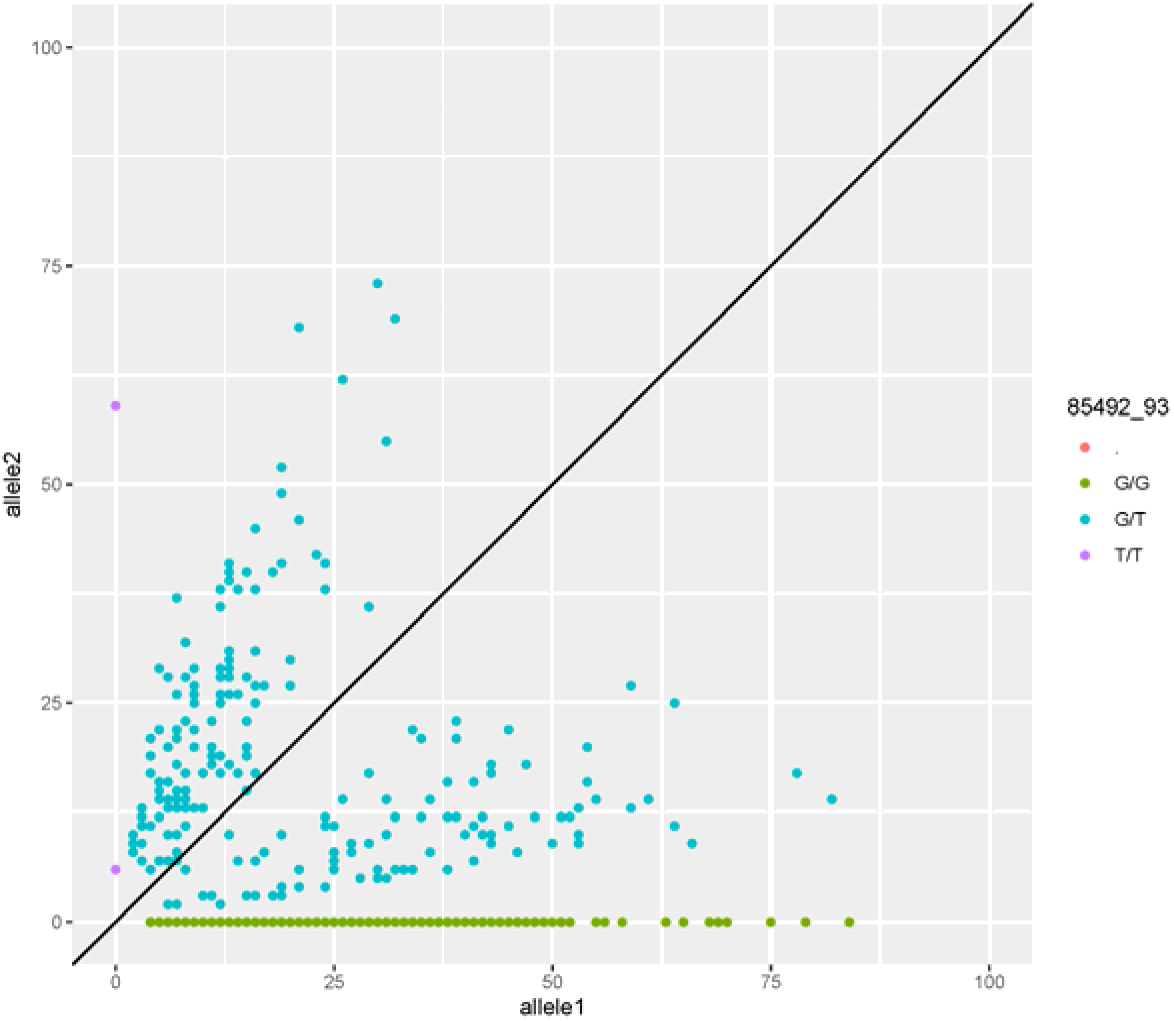
Scatter plot of reads for SNP 85492_93, the SNP on Ots12 associated with the jack life-history. Each dot is a genotype with the reads for the G allele on the x-axis and the reads for the T allele on the y-axis. Genotypes are color coded by the assigned genotype from Stacks. The solid line is the expected ratio of 1:1 for diploid heterozygous genotypes.

## Notes

### Competing Interest Statement

The authors have declared no competing interest.

